# Exocyst Dynamics During Vesicle Tethering and Fusion

**DOI:** 10.1101/354449

**Authors:** Syed Mukhtar Ahmed, Hisayo Nishida-Fukuda, Yuchong Li, W. Hayes McDonald, Claudiu Gradinaru, Ian G. Macara

**Affiliations:** Department of Cell and Developmental Biology, Vanderbilt University School of Medicine, Nashville TN 37240 USA; Department of Physics, University of Toronto, Toronto, ON, M5S 1A7, Canada; Department of Chemical & Physical Sciences, University of Toronto Mississauga, Mississauga, ON, L5L 1C6, Canada; Department of Biochemistry, Vanderbilt University School of Medicine, Nashville, TN 37240, USA

**Author notes:** Equal contributions (Syed Mukhtar Ahmed, Hisayo Nishida-Fukuda). Current address: ^§^Department of Hepato-Biliary-Pancreatic and Breast Surgery, Ehime University, Japan. Corresponding author. Address correspondence to: Ian Macara, Ph.D. Louise B. McGavock Professor, and Chair Dept. of Cell and Developmental Biology Vanderbilt University School of Medicine PMB 407935 U 3209 MRB III Nashville TN 37240-7935 USA.

**Keywords:** vesicles, membrane fusion, dynamics, TIRFM, CRISPR, gene-editing, protein complex

## Abstract

The exocyst is a conserved octameric complex that tethers exocytic vesicles to the plasma membrane prior to fusion. Exocyst assembly and delivery mechanisms remain unclear, especially in mammalian cells. Here we tagged multiple endogenous exocyst subunits with sfGFP or Halo using Cas9 gene editing, to create single and double knock-in lines of mammary epithelial cells, and interrogated exocyst dynamics by high-speed imaging and correlation spectroscopy. We discovered that mammalian exocyst is comprised of tetrameric subcomplexes that, unexpectedly, can associate independently with vesicles and plasma membrane and are in dynamic equilibrium. Membrane arrival times are similar for subunits and vesicles, but with a small delay (~80msec) between subcomplexes. Departure of Sec3 occurs prior to fusion, whereas other subunits depart just after fusion. Single molecule counting indicates ~9 exocyst complexes associated per vesicle. These data reveal the mammalian exocyst as a remarkably dynamic two-part complex and provide important new insights into assembly/disassembly mechanisms.

---

Traffic between membrane-bound compartments requires the docking of cargo vesicles at target membranes, and their subsequent fusion through the interactions of SNARE proteins. The capture and fusion of vesicles are both promoted by molecular tethers known as multisubunit tethering complexes^1^. One group of such tethers, sometimes called CATCHR (complexes associate with tethering containing helical rods) comprises multisubunit complexes required for fusion in the secretory pathway, and includes COG, Dsl1p, GARP and the exocyst^2^. The endolysosomal pathway contains two different tethering complexes, CORVET and HOPS, with similar overall structures to the CATCHR group^3^.

COG consists of two subcomplexes, each containing four subunits, which function together within the Golgi^4-6^. The exocyst is also octameric, and is necessary for exocytic vesicle fusion to the plasma membrane (PM), but the organization of the complex has been controversial^7-10^. Several studies in yeast suggest that one (Sec3) or two (Sec3 and Exo70) subunits associate with the PM and recruit a vesicle-bound subcomplex of the other subunits, but other work argues that the exocyst consists of 2 subcomplexes of 4 subunits each that form a stable octamer or, in mammalian cells, that 5 subunits at the PM recruit 3 other subunits on the vesicle^11-22^. Rab GTPases promote exocyst binding to the vesicle, and SNARES, Rho family GTPases, the PAR3 polarity protein, and phospho-inositide binding domains are all involved in recruiting exocyst to the PM^20, 23-30^.

The dynamics and regulation of assembly and disassembly of these tethering complexes remain unresolved. In mammalian cells the over-expression of individual exocyst subunits causes aggregation and degradation^31^. A pioneering approach to avoid this problem involved silencing the Sec8 subunit and replacement by a Sec8-RFP fusion^31^. Sec8-RFP arrival at the PM was tracked using total internal reflection microscopy (TIRFM), which occurred simultaneously with vesicles ~7.5 sec prior to vesicle fusion^31^. However, the behavior of other exocyst subunits was not addressed. In budding yeast, vesicles remain tethered for about 18 sec prior to fusion, and several exocyst subunits were shown to depart simultaneously at the time of fusion, suggesting that the complex does not disassemble^21^. However, the time resolution was only ~1 sec, so rapid dynamics could not be tracked.

The advent of CRISPR/Cas9-mediated gene editing coupled with the development of high efficiency scientific CMOS (sCMOS) cameras has the potential to revolutionize our understanding of protein dynamics in the living cell. We have exploited these technologies to generate multiple tagged alleles of exocyst subunits by gene editing, and coupled proteomics with high speed TIRFM and fluorescence cross-correlation spectroscopy (FCCS) to quantify exocyst dynamics in unprecedented detail. We discovered that, in mammary epithelial cells, exocyst connectivity is different from previous models of the mammalian exocyst but has similarities to the connectivity in budding yeast^19^. Unexpectedly, each subcomplex can associate with the PM independently of the other, but both are required for vesicle docking. Subunit arrival at the PM coincides with vesicle arrival but with an arrival delay between subcomplexes of ~80 msec. However, one subunit, Sec3, which is part of Subcomplex 1 (SC1), departs prior to fusion and the departure of other subunits, and exhibits anomalous diffusion. Cross-correlation of Sec3 to other subunits is significantly reduced. Taken together, these data are inconsistent with prior exocyst models and suggest that, in mammalian cells, exocyst subunits are in dynamic equilibrium with assembled complexes and the plasma membrane, that intact subcomplexes assemble on secretory vesicles as they dock, and that Sec3 is released prior to fusion.

## RESULTS

### Generation and validation of functional, endogenously tagged exocyst subunits

Each of the 8 exocyst subunits can be C-terminally tagged in *S. cerevisiae* without disrupting function^19^. Therefore, we attempted to incorporate superfolder (sf) GFP, mScarlet-i (Sc), or Halo tags in-frame C-terminally into all exocyst subunit genes of the NMuMG murine mammary epithelial cell line, using Cas9-mediated gene editing (Fig 1a and supplementary Fig 1a). Successfully gene-edited cells were isolated by FACS, and single colonies of cells were expanded and genotyped (Fig 1b and Supplementary Fig 1b-h). GFP+ cell lines were recovered with 5 of the 8 targeting vectors (Fig 1c). For the other subunits, either no GFP+ cells were detected (Exo84), or rare single GFP+ cells were detectable after transfection (Supplementary Fig. 1i-j) but did not proliferate (Sec10, Sec15). Tagging these subunits at the N-terminus was also unsuccessful. NMuMG cells are unusually sensitive to defects in membrane protein delivery, as is caused by silencing of exocyst subunit expression, and rapidly apoptose^11, 24, 30, 32-35^. Therefore, we conclude that tagging Sec10, Sec15 or Exo84 disrupts exocyst function, while C-terminal tags on the other 5 subunits (Exo70, Sec3, Sec5, Sec6, and Sec8) are well tolerated. Homozygous clones of Exo70-GFP, Sec5-GFP, and Sec8-GFP were isolated, plus heterozygous clones of Sec3-GFP and Sec6-GFP (Fig 1c, and Supplementary Fig. 1b-e,g). We also created homozygous double knock-in cell lines of Sec5-Sc or –Halo + Exo70-GFP (Supplementary Fig. 1h).

**Figure 1.**
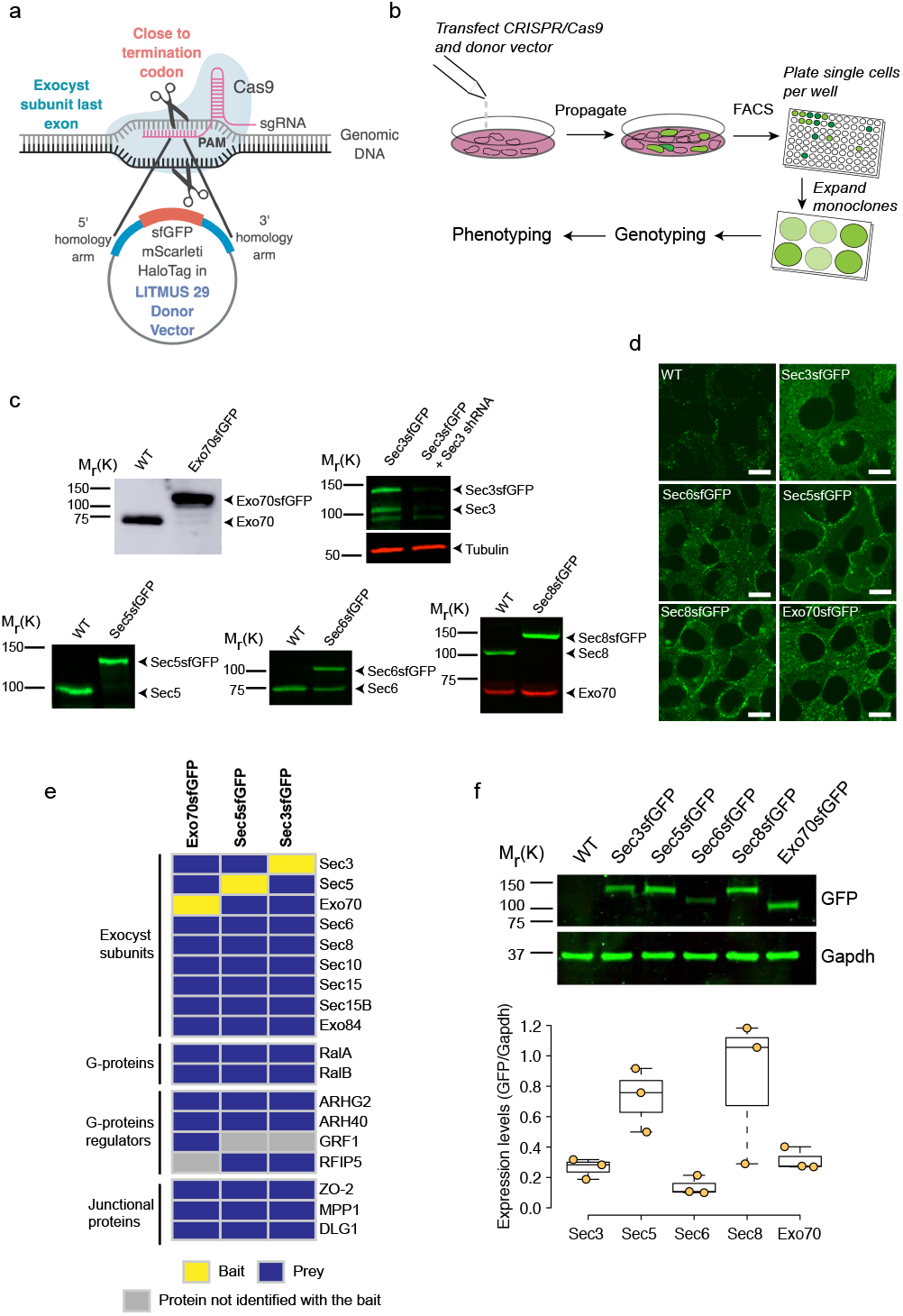
Establishment and Validation of Tagged Exocyst Subunit Cell Lines. (a) Schematic showing region of subunit gene targeted by sgRNA and tags to be inserted C-terminally, in-frame with the coding region. (b) Strategy to isolate CRISPR/Cas9-mediated sfGFP-tagged clones of exocyst subunits in NMuMG cells. (c) Western blots with subunit specific antibodies show successful incorporation of sfGFP in both alleles (Exo70, Sec5 and Sec8), or in one allele (Sec3 and Sec6). A shRNA specific to Sec3 was used to confirm the identity of multiple bands in the blot. (d) Confocal images of live wild-type NMuMG cells and endogenously tagged Sec3-GFP, Sec5-GFP, Sec6-GFP, Sec8-GFP and Exo70-GFP cell lines. Scale bar = 10μm. (e) Protein-protein interaction heat map for endogenous Sec3-GFP, Sec5-GFP and Exo70-GFP pulldown using GFP-Trap nanobodies and MS. Yellow indicate baits, blue the protein identified with high confidence, and grey denotes preys that were not identified with all three baits. The experiments were repeated 3 times for Exo70-GFP and twice for Sec3-GFP and Sec5-GFP. All experiments were also partially confirmed by IP-WB in at least 3 independent experiments, with similar results. (f) Western blot analysis to assess relative abundance of exocyst subunits fused to sfGFP, using anti-GFP antibodies. Gapdh was the loading control. Y-axis shows the ratio of GFP/Gapdh. The blot is a representative image of 3 independent. Quantification shows pooled data from 3 experiments.

Significant GFP fluorescence above the autofluorescence background was detectable by confocal microscopy of live cells for each of the 5 single knock-in lines (Fig. 1d). Sec5, Sec6, Sec8 and Exo70 were enriched at intercellular junctions, consistent with data on fixed and immunostained epithelial cells, but the knock-in cells also have substantial diffuse cytoplasmic pools, and no detectable nuclear signal, contrary to some other reports^36^. Sec3 showed a predominantly cytoplasmic distribution with no significant accumulation at junctions.

To further validate the functionality of the tagged exocyst subunits we performed immunoprecipitations from Exo70-GFP, Sec3-GFP and Sec5-GFP cell lysates using GFP-trap beads, and analyzed them by mass spectrometry (MS). Each tagged protein not only co-precipitated all other 7 subunits (Fig. 1e, and Supplementary Table 1) but also known regulatory factors (RalA and RalB), RhoGEFs, and proteins associated with intercellular junctions ^22, 26, 37-39^. We also identified several peptides from a Sec15-like gene, Sec15L. However, no peptides from 4 Sec6-related genes were detected, suggesting these isoforms are not expressed in NMuMG cells. Independently, Sec8 binding to Exo70-GFP and Sec5-GFP was detected by immunoblots of GFP-trap bead precipitations (Supplementary Fig. 1k).

Interestingly, we also identified the SNARE-associated protein SNAP23 binding to Exo70-GFP, by immunoblot and mass spectrometry of co-immunoprecipitations (Supplementary Fig. 1k, and Supplementary Table 1). However, no SNAP23 association with Sec5-GFP was detectable, suggesting that SNAP23 is likely not associated with the entire exocyst complex. Additionally, we did not detect any Sec1 or other SM proteins. Finally, the tagged cell lines showed similar proliferation rates to one another (Supplementary Fig. 3a). We conclude that the C-terminally-tagged subunits retain normal protein-protein interactions and are biologically functional.

We took advantage of the sfGFP-tag to compare relative expression levels of individual exocyst subunits in NMuMG cells. Surprisingly, as shown in Fig. 1f, these levels are not equimolar. In particular, Sec5 and Sec8 are expressed at super-stoichiometric levels compared to the other 3 subunits (corrected where appropriate for heterozygosity). Sec8-GFP in particular was ~3-fold higher than expected. One interpretation is that addition of the sfGFP-tag stabilizes these proteins. However, blotting of cell lysates for Sec8 from 2 heterozygous Sec8-GFP clones showed that untagged and tagged alleles are expressed at the same levels (Supplementary Fig. 1g). These data suggest that in addition to forming exocyst complexes a surplus of free Sec8 and Sec5 is present in cells. Whether they participate in other protein-protein interactions or have independent functions remains to be investigated.

### Connectivity of the mammalian exocyst

It has been proposed that exocyst in budding yeast is a stable octamer comprised of two subcomplexes (SC1: Sec3, Sec5, Sec6, Sec8 and SC2: Exo70, Exo84, Sec10, Sec15)^19, 40^. To investigate the subunit connectivity in mammalian exocyst we silenced Sec8 in Exo70-GFP cells using a previously validated hairpin RNA^30^, then captured the complex on GFP-trap beads and performed quantitative full scan LC-MS/MS. Experiments were performed 3d post-transduction of shRNAs, before apoptosis occurs, and loss of Sec8 was confirmed by western blot analysis. Interestingly, only proteins corresponding to SC2 were recovered; those from SC1 were absent (Fig. 2a and Supplementary Fig. 2a,b). Loss of Sec8 also strongly reduced detection of RalA and RalB, which bind not only to Sec5 in SC1 but also to Exo84 in SC2 (Supplementary Fig. 2b).

**Figure 2.**
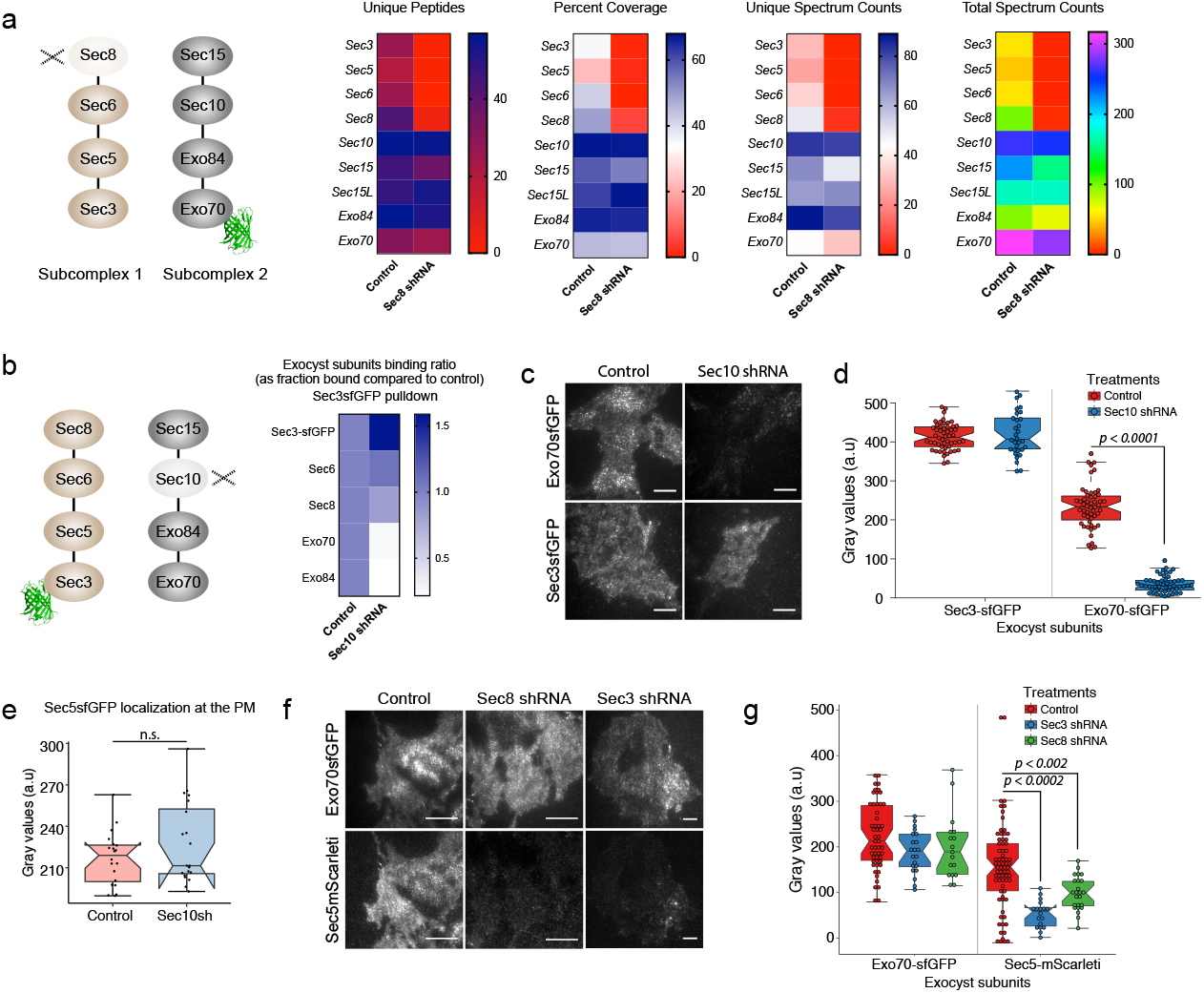
Mammalian Exocyst Complex Comprises Two Subcomplexes that Can Localize to the PM Independently. (a) Quantitative full scan LC-MS/MS analysis of endogenous Exo70-GFP pull-downs from intact or Sec8-depleted cells. Inputs for the samples were calibrated and normalized using MRM-MS analysis on the same samples. Schematic shows the experimental design in which Exo70-GFP was captured using GFP-Trap beads and Sec8 was depleted by shRNA. (b) Heat map summarizing relative binding of Sec6, Sec8, Exo70, and Exo84 to Sec3-GFP in Sec10-depleted cells compared to control shRNA treated cells, as assessed by western blot. Also see Figure S2H-I. (c) TIRFM images of Exo70-GFP and Sec3-GFP in Control or Sec10 shRNA treated cells. Scale bars 20μm. (d) Quantification of relative fluorescence intensities in C. (e) Quantification of fluorescence intensities from TIRFM images of Sec5sfGFP cells treated with Control or Sec10 shRNA. (f) TIRFM images of double knock-in NMuMG cells, showing localizations of Exo70-GFP and Sec5-mScarleti at the PM in Control, Sec8 shRNA and Sec3 shRNA treated cells. Scale bars = 20μm. (g) Quantification of relative fluorescence intensities in D. Center lines show the median; box limits indicate 25^th^ and 75^th^ percentiles; whiskers extend 1.5X the IQR from the 25^th^ and 75^th^ percentiles. Experiments were repeated at least 3 times with similar results.

Conversely, we silenced Sec10 expression^30^ by RNAi and captured Sec3GFP then subjected the precipitate to immunoblotting for Sec6, Sec8, Exo70 and Exo84 (Fig. 2b and Supplementary Fig. 2h-i). In this experiment components of SC1 (Sec6, Sec8) coprecipitated with Sec3, but not those of SC2 (Exo70, Exo84). Together, these data argue that the mammalian exocyst connectivity is similar to that in yeast, with 2 subcomplexes, each of which needs to be intact for interaction with the other, and that loss of one subunit disrupts the stability only of its cognate subcomplex.

The functionality of the subcomplexes has never been addressed. To explore this issue we asked if they can associate with the PM independently or only as an octameric complex. Association with basal PM was assessed by TIRFM. Subconfluent cells were used because vesicle fusion frequency at the basal membrane decreased substantially with confluency, when vesicles are redirected to apical junctions^41^. Sec3- and Sec8-GFP (SC1) and Exo70-GFP (SC2) were detected as fluorescent puncta in the unperturbed cell lines, with ~70% Exo70-GFP and Sec8-GFP associated with vesicles (Supplementary Fig. 2l-n). Unexpectedly, however, when Sec10 (in SC2) was silenced Sec3-GFP remained detectable on the basal membrane, although Exo70-GFP was lost (Fig. 2c-d). Sec8 association with vesicles was slightly reduced (to ~45%; Supplementary Fig. 2m,n). A similar experiment using Sec5-GFP cells showed no impact of silencing Sec10 on the association of Sec5 with the membrane (Fig. 2e).

Conversely, silencing of Sec8 or Sec3 (in SC1; Supplementary Fig. 2c) caused depletion of Sec5 (SC1) but not Exo70 (SC2) from the membrane (Fig. 2f-g, and Supplementary Fig. 2j). Consistent results were also obtained when Exo70 was deleted by Cas9-mediated gene editing: Sec8 (in SC1) was retained at the membrane (Supplementary Fig. 2f-g,k). From these data we conclude that each subcomplex remains intact in the absence of the other subcomplex, and, unexpectedly, retains association with both the PM and with vesicles.

### Exocyst dynamics during vesicle delivery and fusion

Do all exocyst subunits arrive at the PM together with an exocytic vesicle, or do the subcomplexes or individual subunits arrive separately? To address this question we transfected NMuMG cells with vectors to express mApple-Rab11 as a vesicle marker plus VAMP2-pHluorin, a vesicle-targeted pH-sensitive GFP that becomes highly fluorescent upon vesicle fusion with the PM. Arrival of Rab11+ vesicles at the membrane can then be correlated with the fusion event by 2-channel time-lapse TIRFM (Fig. 3a). A typical kymograph is shown in Fig. 3b, and analysis of multiple events provided a median value of −14.5 sec for arrival (Fig. 3c).

**Figure 3.**
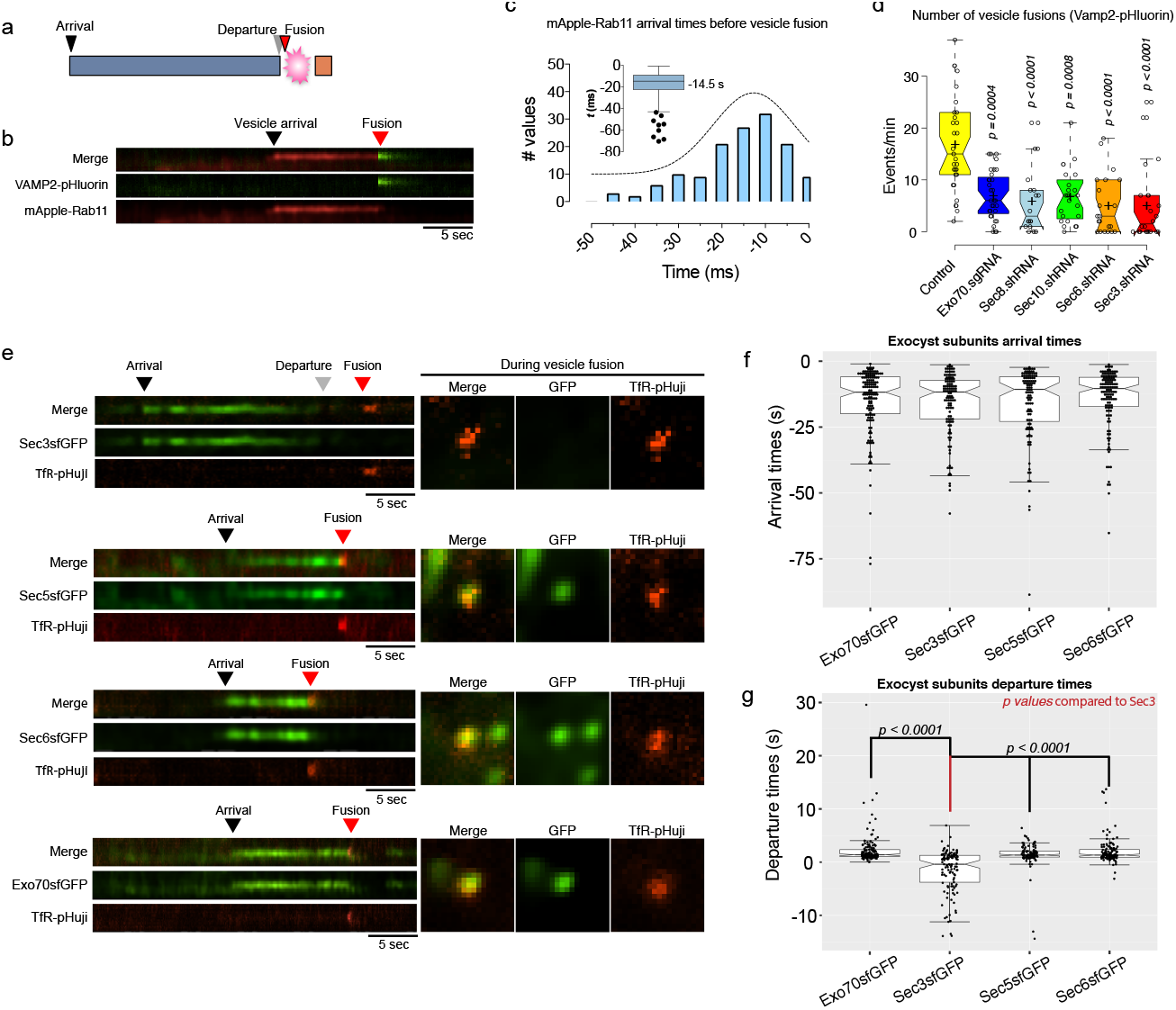
Sec3 Departs from the Exocyst Complex Prior to Vesicle Fusion. (a) Schematic representation of the data. Black arrowhead shows arrival of the target protein, gray arrowhead shows departure, red arrowhead shows vesicle fusion events as demarcated by pH sensitive fluorophores, pHluorin or pHuji, fused to Vamp2 or Transferrin receptor (TfR), respectively. (b) NMuMG cells were transduced with mApple-Rab11 and Vamp2-pHluorin lentivirus and imaged by TIRFM. Kymographs show arrival and departure of mApple-Rab11 and vesicle fusion, highlighted by Vamp2-pHluorin flashes. Speed = 5Hz. Scale bar = 5s. (c) Distribution of mApple-Rab11 arrival times prior to vesicle fusions. Inset shows box and whisker plot; center line = −14.6s (median; 95% CI: −16.6 to −12.5). Error bars = Tukey’s range. (d) Effects of exocyst subunits depletion on vesicle fusion activities in NMuMG cells assessed using Vamp2-pHluorin. n = 32, 32, 18, 22, 21, 22. *P* values compared to control. (e) Kymographs showing itinerary of Sec3-GFP, Sec5-GFP, Sec6-GFP and Exo70-GFP arrivals/departures with respect to the vesicle fusion marker TfR-pHuji. Square images show snapshots in the green and red channels at the time of vesicle fusions. Scale bar = 5s. (f-g) Quantification of exocyst subunit arrival and departure times at and from the vesicle fusion site shown in E. Data points shown as dots. (f) Median arrival times in secs: −11.7 (95% CI: −12.9 to - 9.1; Exo70), −11.6 (95% CI: −14.6 to −10.2; Sec3), −10.6 (95% CI: −14.0 to −8.2; Sec5) and −10.3 (95% CI: −12.3 to −8.8; Sec6). n = 146, 132, 108, and 132 objects in the order data shown. (g) Median departure times in secs: 1.4 (95% CI: 1.2 to 1.5; Exo70), −0.4 (95% CI: −0.98 to −0.69; Sec3), 1.3 (95% CI: 1.2 to 1.6; Sec5) and 1.3 (95% CI: 1.2 to 1.7; Sec6). n = 145, 117, 108, and 132 particles from 33, 19, 29, and 26 cells in the order data are shown. Center lines = medians; hinges extend from 25^th^ and 75^th^ percentiles. Statistical significance measured by Kruskal-Wallis test followed by Dunn’s *post hoc* analysis. Experiments were repeated 3 times with similar results.

We next asked if vesicle fusion requires exocyst, from measurements of events/min/field in cells separately depleted of each of 5 subunits (Supplementary Fig. 2c-g). Because silencing of Exo70 by shRNA was not efficient we used gRNA targeted gene disruption for this subunit (Supplementary Fig. 2j). Importantly, in each case, loss of expression of subunits from either SC1 or SC2 caused a significant reduction in fusion events (Fig. 3d). We conclude that although subcomplexes are competent to associate with the PM, they cannot separately promote vesicle fusion.

Based on this foundation, we simultaneously tracked exocyst subunit arrival and vesicle fusion using our GFP knock-in cell lines transfected with transferrin receptor fused to pHuji, a pH-sensitive RFP (TfR-pHuji). Representative kymographs are shown in Fig. 3e, and data are quantified in Fig. 3f. Interestingly, the 4 tested subunits (Sec3, Sec5, Sec6, Exo70) all arrive with median times of 10 - 14 sec prior to fusion, which is close to the arrival time of the Rab11+ vesicles. This concordance is similar to that observed in budding yeast^21^, and suggests that exocyst arrives at the PM with a vesicle, rather than being pre-assembled on the membrane.

We next asked when exocyst subunits disappear from fusion sites. Sec5, Sec6 and Exo70 each depart at a median time of ~1.3 sec post-fusion. Unexpectedly, however, (Fig. 3g) Sec3 was anomalous with a median departure time 0.4 sec prior to fusion, significantly different from the other subunits. This result is consistent with the reduced enrichment of Sec3-GFP detected at intercellular junctions (Fig. 1d), and implicates Sec3 in a unique function associated with vesicle fusion. An alternative interpretation is that Sec3-GFP fusion might be only partially functional, with defective binding to the exocyst complex. We do not believe this is the case, however, because the Sec3-GFP cell line proliferates normally (Supplementary Fig. 3a), and the abundance of Sec5-Halo at the PM is identical in lines that express Sec3-GFP versus Sec8-GFP (Supplementary Fig. 3b). Furthermore, we did not observe any differences in subunit interactions whether we used heterozygous Sec3-GFP or a homozygous Sec3-Sc (Supplementary Fig. 3c,d). However, our ability to employ Sec3-Sc for imaging was limited by rapid photobleaching of this fluorophore.

### Coincidence measurements between exocyst subunits by TIRF

To test whether subunit arrivals at the PM are truly coincident we used double knock-in cell lines Sec8-sfGFP+Sec5-Halo or Exo70-sfGFP+Sec5-Halo and assessed their fluorescence intensity trajectories with high time resolution. Sec8-sfGFP and Sec5-Halo almost always arrived in the TIRF field together; strikingly, however, Exo70-sfGFP showed a median delay of 77ms with respect to Sec5-Halo (Fig. 4a,b). Importantly, this small but significant lag between SC1 and SC2 subunits could not have been detected by comparing single subunit versus vesicle fusion arrival times, as described in Fig. 3 and in other studies^21^.

**Figure 4.**
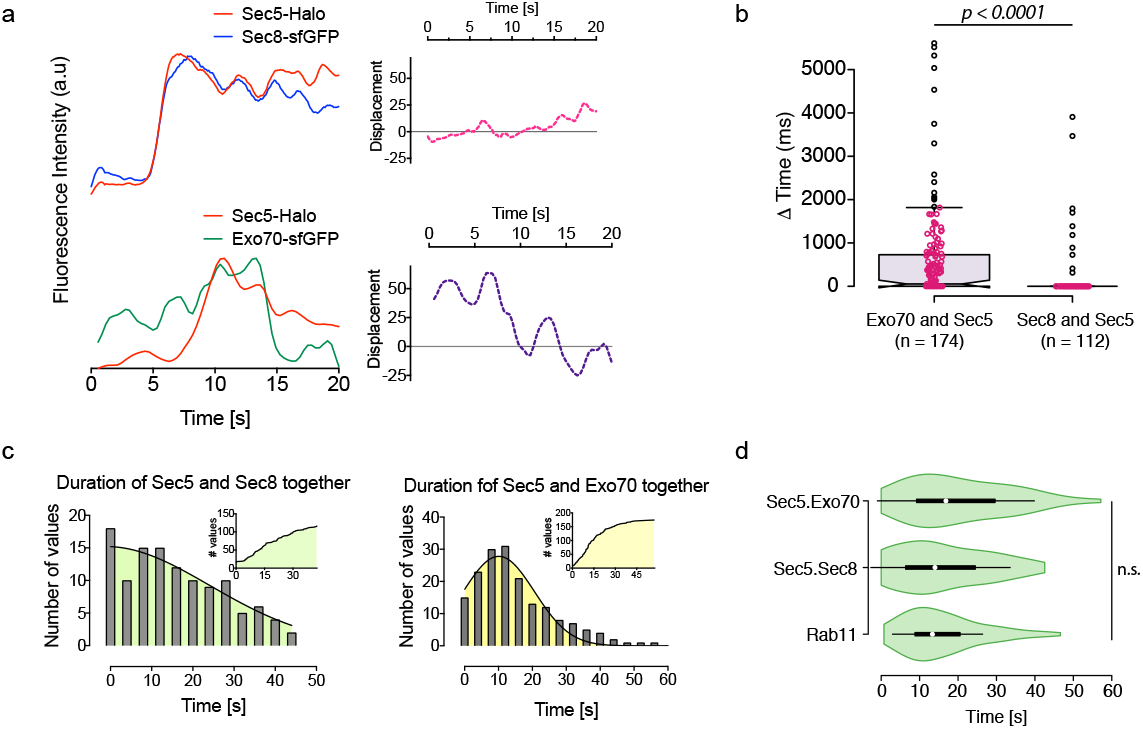
Coincidence Measurements Between Exocyst Subunit Pairs in the Same Subcomplex or Different Subcomplexes by TIRF in Live Cells. (a) Fluorescence intensity trajectories (intensity values were normalized to a common scale) for the Sec5-Halo+JF585/Sec8-GFP pair, and Sec5-Halo+JF585/Exo70-GFP pair over 20 sec. The fluorophores were excited simultaneously and images captured at 12.5Hz (left). On the right, graphs show the displacement of the normalized intensities of the two fluorophores at each time point. (b) Quantification of the delay in coincidence between Exo70-GFP and Sec5-Halo+JF585 or Sec8-GFP and Sec5-Halo, measured by TIRFM. Outliers are represented by black circles; data points are plotted as magenta circles. The *n* denotes number of particles analyzed from a total of three experiments. (c) Distributions of the duration of the indicated subunits were together in the TIRF field. (d) Comparison on the distribution of residence times of Rab11 and the indicated pairs of exocyst subunits. White circles = medians; box limits indicate the 25^th^-75^th^ percentiles; whiskers extend 1.5X the interquartile ranges from 25^th^-75^th^ percentiles; polygons represent density estimates of data and extend to extreme values. Statistical significance measured by Kruskal-Wallis test followed by Dunn’s *post hoc* analysis.

We also measured residency times for exocyst subunits on the PM. Both Exo70-sfGFP + Sec5-Halo and Sec8-sfGFP + Sec5-Halo pairs could be traced together for ~12-14s (Fig. 4c), and these durations were similar to that of vesicle residency at the PM as assessed using Rab11-mApple (Fig. 3c,4d). These observations indicate that while association between SC1 subunits Sec5 and Sec8 is stable and they arrive together at the membrane, association between SC1 and SC2 is dynamic and they can arrive independently at (or near) the PM. Nevertheless, once at the PM SC1 and 2 remain associated, with a residence time similar to that of the vesicles.

### Single molecule approach to quantifying exocyst subunit interactions

Whether vesicles destined for the PM are associated with intact exocyst or with only a subset of exocyst subunits remains controversial. To measure protein-protein interactions between subunit pairs throughout the cytoplasm, rather than just at the PM, we used single molecule counting.

First, we needed to determine the stoichiometry of dye binding to Halo, as this parameter has not been previously reported. To do so, we created a YFP-Halo fusion protein, expressed it in NMuMG cells (Supplementary Fig. 5a) and added Halo dye (J585-HTL) at different concentrations for various periods. Flow cytometry showed a strong linear correlation between YFP and dye fluorescence (Supplementary Fig. 5b). Next, cells were ruptured in a small volume, which was cleared by centrifugation, and supernatants were spread onto coverglasses for single particle counting by two-color TIRFM (Fig. 5a). Maximum fractional labeling efficiency was 0.40 ± 0.014 after 1.5 hrs, and increasing incubations to 18 hrs did not increase fractional labeling beyond 0.4 (Supplementary Fig. 5c,d). Additionally, FCCS measurements robustly detected a value of 0.4 for cross correlation between Halo and YFP fluorescence in intact cells (Supplementary Fig. 5e-h). Therefore, this value was used to correct our data on Halo-tagged subunit interactions.

**Figure 5.**
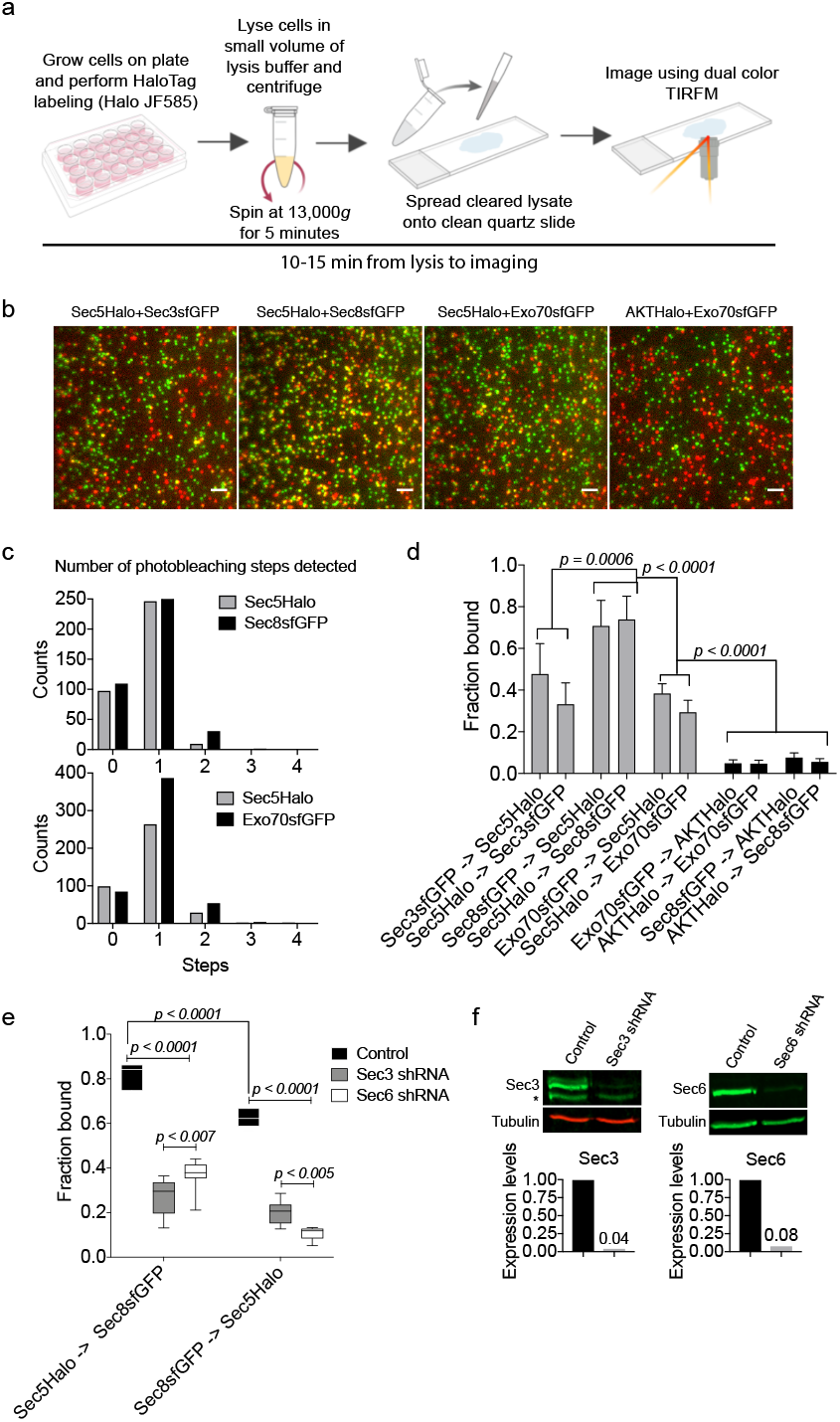
Assessment of Fractional Binding of Exocyst Subunits Using Single Molecule Approach. (a) Diagram showing experimental workflow. (b) Cleared cell lysates from the indicated double knock-in cells were spread between quartz slides and glass coverslips for TIRF imaging. The GFP and Halo+JF585 fluorophores were excited simultaneously with 488nm and 561nm lasers at 12.5Hz for 40s. Scale bars = 1μm. (c) Number of photobleaching steps detected for Sec5-Halo and Sec8-GFP in experiments shown in panel B. (d) Quantification of the fraction of subunit “A” bound to (->) subunit “B” from experiments shown in panel B. Numbers of particles analyzed for each pair are shown in Supplementary Fig. 4c. (e) Fraction of Sec5-Halo and Sec8-GFP bound to each other in Sec3 or Sec6 shRNA treated cells compared to control. Center line denotes mean. (f) Western blot analysis of Sec3 and Sec6 shRNA knockdown efficiencies in the experiment shown in E. Error bars denote ± *s.d.* Statistical significance was computed by one-way (D) or two-way (E) ANOVA, followed by Tukey’s multiple comparison tests. Experiments were repeated 3 times and data were pooled.

Next, lysates from double knock-in cells expressing Sec5-Halo+Sec3-GFP, Sec8-GFP, or Exo70-GFP were rapidly spread on coverslips and co-localization in single particles was quantified as described above (Fig. 5b). The majority of detected spots contained only single molecules of each species, as assessed by photobleaching, thus enabling a reliable assessment of fractional inter-subunit binding (Fig. 5c, Supplementary Fig. 4d). Sec3 counts were corrected for heterozygosity. An AKT-Halo fusion, expressed in cells at a similar concentration to the endogenous Exo70-GFP fusion (Supplementary Fig. 4a,b), was used as a negative control to correct for chance colocalization. At the fluorophore concentrations present, overlap was only 5 – 8% (Fig. 5d).

Coincidence within the diffraction limit between Sec8-GFP and Sec5-Halo was much higher, at ~70% (Fig. 5d, Supplementary Fig. 4c). Surprisingly, however, only ~40% of Sec3- and Exo70-GFP molecules colocalized with Sec5-Halo (Fig. 5d). Conversely, colocalization of Sec5-Halo to Sec3-GFP or Exo70-GFP was even lower, at ~30%, likely because Sec5 is expressed at a significantly higher level than Sec3 or Exo70 (Fig. 1f). These results were corroborated by co-immunoprecipitation of Sec8GFP with ~80% of total Sec6 (in SC1) but only ~30% of total Exo70 (in SC2) (Supplementary Fig. 4e,f). Together, our data suggest that some subunits within a subcomplex (Sec5, Sec6 and Sec8 in SC1) interact stably, but that Sec3 binding is substoichiometric, and similar to the weak interaction between subcomplexes.

Because Sec3 binds to other SC1 subunits so weakly, we asked if it is required for subcomplex stability. Sec3 or Sec6 (as positive control) were efficiently silenced and Sec5/Sec8 interactions were measured by single molecule counting. Surprisingly, both perturbations strongly reduced Sec5/Sec8 binding (Fig. 5e,f). These data argue that Sec3 is essential for the assembly of SC1 (and perhaps interaction with SC2), but not for maintenance of the Sec5/6/8 heterotrimer once it has formed. This idea is consistent with the early departure of Sec3 from the exocyst complex prior to vesicle fusion (Fig. 3e,g)

### Fluorescence cross-correlation spectroscopy of exocyst subunits suggests a fraction of the complex is pre-assembled

As an independent approach to determine the fractional association of SC1 and SC2 we performed in-cell dual-color FCCS. We assessed co-diffusing fractions for Exo70-GFP, Sec8-GFP or Sec3-GFP with Sec5-Halo labeled with JF-646-Haloligand (Eq 3). About 70% of Sec8-GFP was coupled to Sec5-Halo within the cell and 64% near the bottom of the cell (Fig. 6a,d; Supplementary Fig. 6a,d), an abundance similar to that assessed by single molecule counting (Fig. 5d). Sec3-GFP/Sec5-Halo coupling was ~50% within the cell and ~44% near the bottom (corrected for heterozygosity) (Fig. 6b,d; Supplementary Fig. 6b,d). We also assessed the association between SC1 and SC2 from the Sec5-Halo/Exo70-GFP pair, with ~48% of the cytosolic fractions co-diffusing (Fig. 6c). The fraction co-diffusing near the bottom membrane was ~40% (Supplementary Fig. 6c,d), but this is likely an underestimate because of contributions from both membrane and cytoplasm. To address this more rigorously we measured co-localization within the TIRF field, which showed ~85% of Sec5-Sec8 and ~60% of Sec5-Exo70 subunits are together (Fig. 6e).

**Figure 6.**
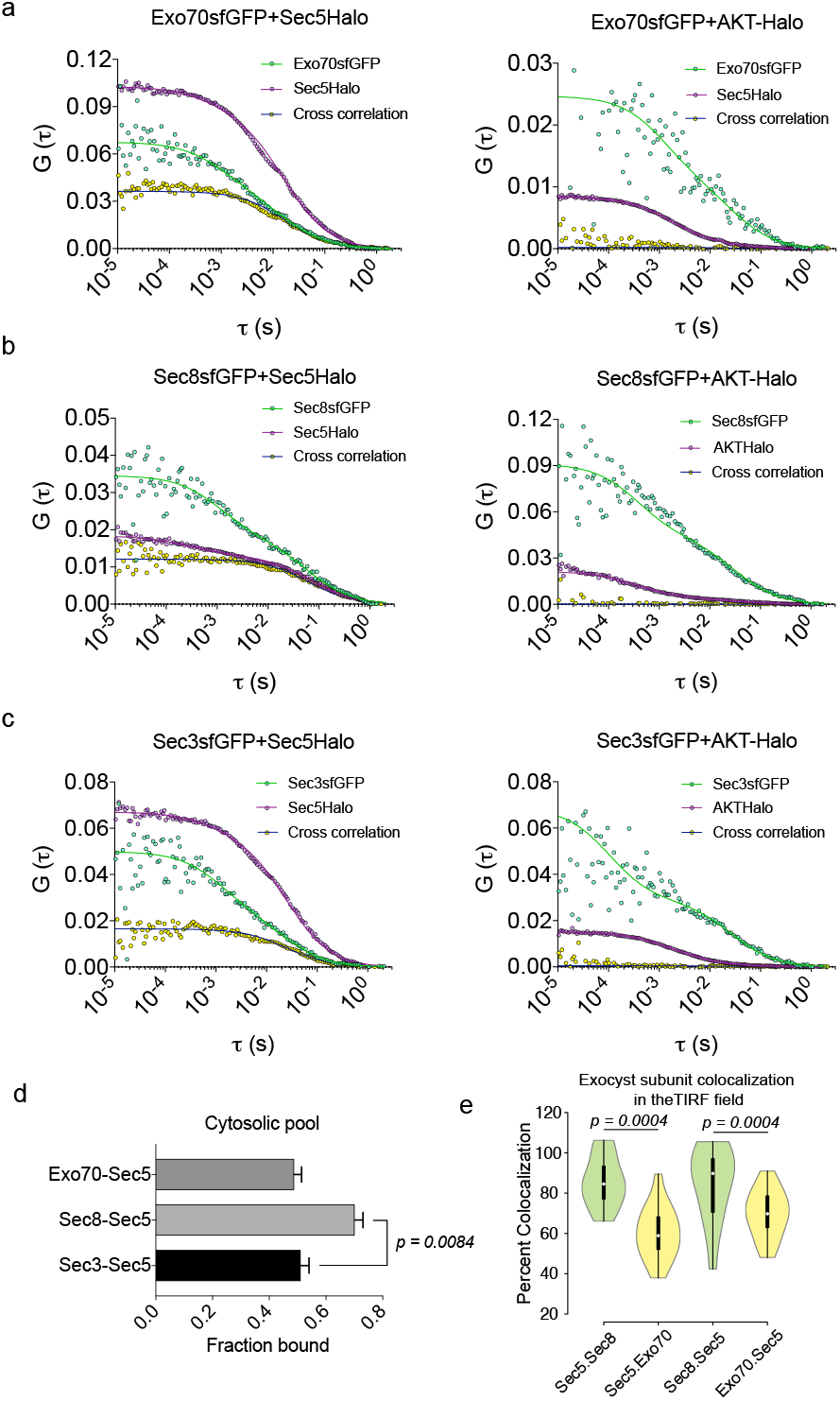
Dual-color FCCS of Exocyst Subunits in Membrane and Cytosol. (a) Sec5-Halo and Exo70-GFP FCS measurements in the cytosol and the PM. Exo70-GFP and AKT-Halo fluorescence cross-correlation was used as negative control. HaloTag was labeled using JF646 Halo ligand (200 nM for 1.5h). (b) Sec5-Halo and Sec8-GFP, or Sec8-GFP and AKT-Halo FCS measurements in the cytosol or the PM. (c) Sec5-Halo and Sec3-GFP, or Sec3-GFP and AKT-Halo FCS measurements in the cytosol or the PM. (d) Statistics of fraction of GFP-tagged exocyst subunit bound in the cytosol. Sec8-GFP+Sec5-Halo: 70% ± 2.9%, Sec3-GFP+Sec5-Halo: 51% ± 2.9%, Exo70-GFP+Sec5-Halo: 48% ± 2.6% (mean ± *s.e.m.).* (e) Fractional coincidence between Sec5-Halo and Sec8-GFP or Sec5-Halo and Exo70-GFP was analyzed using TIRFM. Calculations were corrected for HaloTag labeling efficiency and expression levels of Sec8-GFP and Exo70-GFP with respect to Sec5-GFP. Centerline denote median. Sec5-Halo bound to Sec8-GFP was 85% ± 3.3% (mean ± *s.e.m).* Sec8-GFP bound to Sec5-Halo was 85% ± 5.1%. Sec5-Halo bound to Exo70-GFP was 60% ± 3.2% and Exo70-GFP bound to Sec5-Halo was 69.8% ± 2.9%. Object detection parameters were set between 0.30μm and 0.35 μm and contrast adjusted to faithfully detect objects of interest. White circles show the medians; box limits indicate the 25th and 75th percentiles as determined by R software; whiskers extend 1.5 times the interquartile range from the 25th and 75th percentiles; polygons represent density estimates of data and extend to extreme values. *p-values* were calculated using Kruskal-Wallis test followed by Dunn’s multiple comparison test. The hydrodynamic radius (R_H_) for the slow diffusing fraction was median 466nm (95% CI: 355.2 to 606.8), and that of the fast diffusing component, 14.82nm (95% CI: 13.22 to 17.27). FCS measurements (dots = data points) were fitted (lines) to the model described in Methods (Eqs 1 and 2). Curves shown are representative, taken from a single cell. Statistics are from >25 measurements for each condition from 2 experiments and >5 cells. *p values* were calculated by one-way ANOVA followed by Tukey’s *Post hoc* test. R_H_ was calculated using the Stokes-Einstein equation (Eq 4).

Data fitting (Eqs 1 and 2), revealed two diffusion components, with the slower (49% ± 10%) GFP-tagged population (*D* = 0.54μm^2^/s ± 0.26) within the cell being common among all the subunits and cross-correlating. The faster component, which does not crosscorrelate, has a median diffusion rate of 14.5μm^2^/s, reminiscent of freely diffusing molecules in cells^42^. Importantly, Exo70-GFP, Sec8-GFP and Sec3-GFP did not show any crosscorrelation with the negative control, Akt-Halo (Fig. 6a-c). From these experiments we conclude that about half of all exocyst subunits in the cells are likely associated with vesicles, while the remaining subunits are freely diffusing monomers and (sub)complexes.

### Diffusivity of exocyst subunits at the plasma membrane

The weak association of Sec3-GFP with intercellular junctions (Fig. 1d) suggests that this subunit has unique properties. Localization of Sec3-GFP at the plasma membrane is also significantly lower than that of its subcomplex partners (Fig. 1d, 7a-b). These results, together with the observation that Sec3-GFP leaves the docked complex before the other subunits (Fig. 3g), suggest that Sec3 interacts unusually weakly with the complex. To further test this possibility, we measured subunit diffusivity at the PM by tracking individual particle trajectories, and calculated their mean squared displacements (Fig. 7c). Sec3-GFP diffusion was anomalous with a median 0.12μm^2^s^-1^ MSD compared to other subunits, which ranged from medians of 0.02μm^2^s^-1^ to 0.08 μm^2^s^-1^ (Fig. 7d-e). Total displacement measurements showed that Sec3-GFP motion was higher than that of other subunits, with a median of 0.21 μm as compared to Sec8-GFP or Exo70-GFP (median 0.10μm), and Sec5-Halo (median 0.07μm) (Supplementary Fig. 7a). We conclude that Sec3 interacts with the complex with unusually low affinity, suggesting that it might be the limiting factor in formation of octamer.

**Figure 7.**
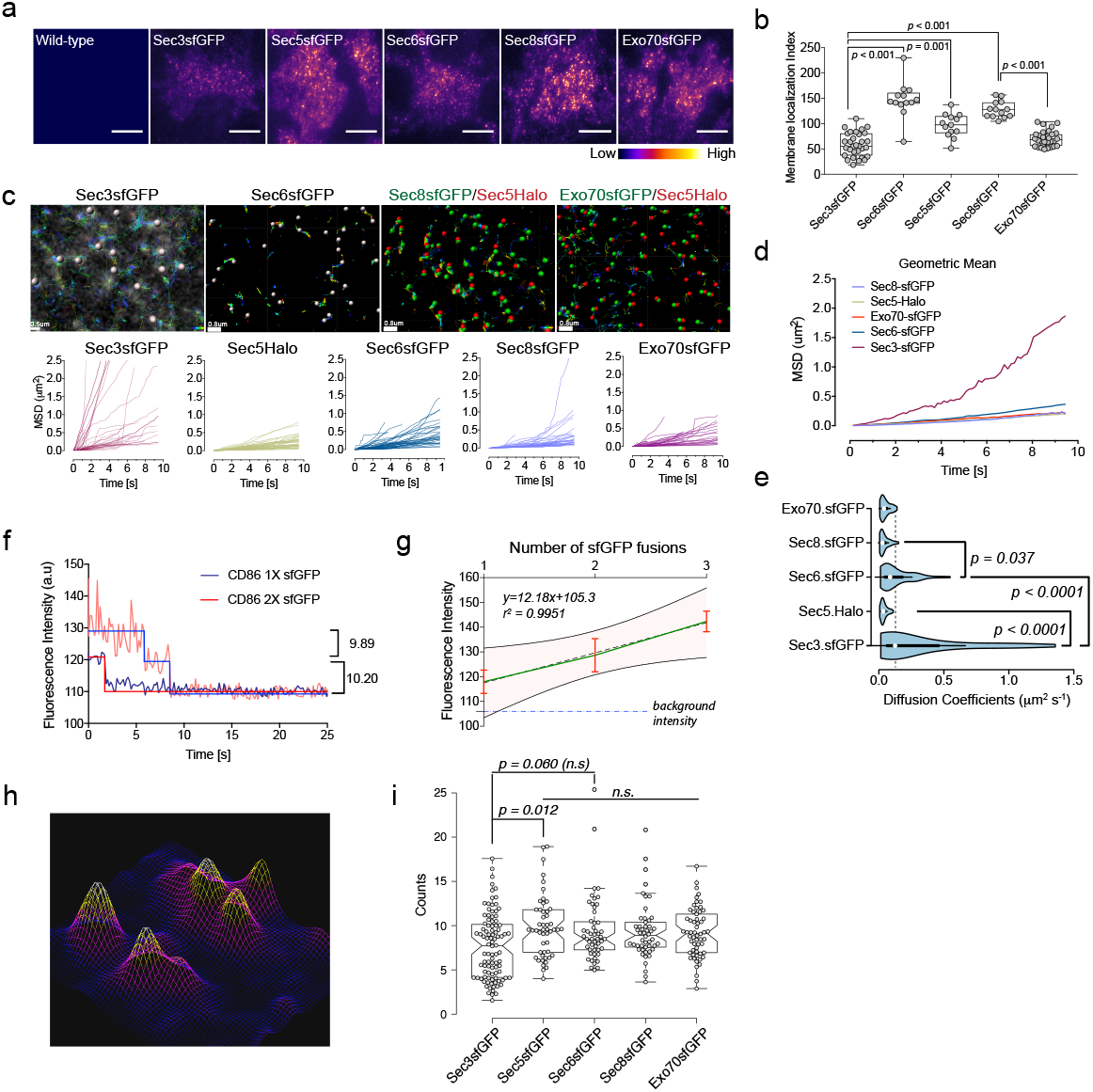
Measurements of Diffusivity of Exocyst Subunits. (a) TIRFM images of untagged wild-type or indicated exocyst subunits fused to sfGFP (top). Heat map of above images (bottom). Scale bar = 20μm. (b) Quantification of exocyst subunits localization at the TIRF field. Membrane localization Index = Density of spots * Intensity of cells. Whiskers extend from min to max. Values for Sec3 and Sec6 were corrected for heterozygosity. n= 31, 13, 13, 13, 32 fields containing 2-4 cells each. (c) Particle tracking over time for the subunits indicated. For two color tracking, each channel was tracked using Imaris software tracking algorithm and overlaid. Graphs show mean squared displacements over time. Scale bar = 0.5μm (Sec3GFP) and 0.8μm (rest). (d) Representation of data in B as geometric means of MSDs <r^2^>. (e) Diffusion coefficients of the indicated subunits were measured from <r^2^>. Mean (μm^2^ s^-1^) ± *s.e.m.* = 0.31 ± 0.05 (Sec3-GFP), 0.04 ± 0.002 (Sec5-Halo), 0.13 ± 0.01 (Sec6-GFP), 0.04 ± 0.004 (Sec8-GFP), 0.05 ± 0.004 (Exo70-GFP). White circles show the medians; box limits indicate the 25th and 75th percentiles; whiskers extend 1.5 times the interquartile range from the 25th and 75th percentiles; polygons represent density estimates of data and extend to extreme values. (f) Stepwise photobleaching of CD86 fused to one or two GFP molecules used to determine intensities of known numbers of GFP molecules. (g) Standard curve for counting protein molecules, from intensities measured on CD86 fused to 1, 2 or 3 GFP molecules. Error bars = *± s.d.;* pink shaded region denote 95% confidence band. (h) Fluorescence landscape of Sec8sfGFP shows a typical resolvable distribution of molecules within an ROI from which intensity measurements were taken. (i) Exocyst molecule counting at vesicle fusion sites. Intensity (y) was measured from the peak and converted to the number of molecules using the regression equation determined in G. Numbers for Sec3 and Sec6 were corrected for heterozygosity. Mean ± *s.d.* = 7.6 ± 3.6 (Sec3-GFP), 9.8 ± 3.5 (Sec5-GFP), 9.8 ± 3.4 (Sec6-GFP), 9.35 ±3.2 (Sec8-GFP), 9.19 ± 2.8 (Exo70-GFP). Centers indicate median, box limits indicate 25^th^ and 75^th^ percentiles and whiskers extend 1.5x IQR from 25^th^ and 75^th^ percentiles. n = 96, 50, 51, 50, 58. Coefficient of variation (CV) = 47.9%, 35.2%, 39.2%, 34.3, 30.7% in the order indicated in the graph. CV between each subunit was 7.9%. Experiments were repeated at least 3 times with similar results. *P* values were computed using Kruskal-Wallis test followed by Dunn’s multiple comparison tests. All experiments were repeated at least 3 times and results were pooled.

### Quantification of exocyst molecules present during vesicle tethering

To better understand how exocyst tethers vesicles to the membrane, we quantified the numbers of complexes at tethered vesicles before fusion. Stepwise bleaching was not detected, suggesting large numbers of molecules were present, so we used ratio comparison to fluorescent standards^43^. We fused 1, 2 or 3 sfGFPs in tandem to the C-terminus of CD86, a monomeric, PM-localized protein^44^ and determined their photobleaching steps and initial intensities (*l*_o_). These values were then used to generate a standard curve with coefficient of determination >0.99 (Fig. 7f-g). To further ensure that we were measuring monomeric CD86, we immunostained fixed cells with anti-GFP antibodies conjugated to biotin and probed for photobleaching steps using streptavidin-ATTO488 (Supplementary Fig. 7b,d). These experiments confirmed that our constructs behave as expected. We also empirically determined the PFS of our objective and limited our analysis to objects that were resolved by a distance of >4 pixels on the camera to measure peak intensities (Supplementary Fig. 7b-i).

To calculate the numbers of subunits at sites of vesicle fusion, we expressed the TfR-pHuji construct in the exocyst-GFP knock-in cell lines and measured the peak intensities of well resolved particles of exocyst subunits at fusion sites (Fig. 7h-i). We estimate ~9 molecules of each exocyst subunit associate with each vesicle during tethering (Fig. 7i), slightly less than that predicted for budding yeast^45^.

## DISCUSSION

A major goal of cell biology is to quantitatively assess the dynamics of molecular processes in living cells. This goal has often been confounded by perturbations introduced through over-expression of gene products, and by imaging limitations. In this study, we report the first quantitative analysis of endogenous exocyst dynamics in mammalian cells, at unprecedented resolution. We used CRISPR/Cas9-mediated gene editing to incorporate fluorescent tags at the C-termini of 5 exocyst subunits in NMuMG cells and assessed connectivity and abundance. We employed fast TIRF imaging of these endogenous molecules to unravel arrival and departure times at vesicle fusion sites, and single-molecule approaches to estimate the fractional binding of exocyst subunits, and the number of subunit molecules per tethered vesicle. Together, these approaches enable a single cell biochemistry that is broadly applicable, and in this study has provided new and comprehensive insights into mammalian exocyst behavior.

Previously, we reported that Par3 interacts with exocyst in NMuMG cells and is required for membrane protein recruitment to lateral membranes and tight junctions^46^. However, Par3 is absent from the basal membrane, and is unlikely to be involved in recruitment to this region of the cell cortex. Indeed, when we expressed mApple-Par3 in the knock-in cell lines, no signal was detectable at the basal surface by TIRFM; and silencing of Par3 expression did not reduce vesicle fusion frequency there (unpublished observations). Therefore, Par3 is not necessary for basal vesicle tethering and fusion, and whether there are different exocyst receptors for different regions of the cell remains to be investigated.

The exocyst assembly mechanism remains controversial. In yeast, several lines of evidence suggest that exocyst is a stable octameric complex, comprised of 2 subcomplexes, that is bound to vesicles prior to their arrival at the PM^19, 21^. Others have proposed that Sec3 or Exo70 functions exclusively at the PM, and that the remaining exocyst subunits arrive on secretory vesicles^17, 18^. In mammalian cells 5 subunits on the PM were proposed to interact with 3 subunits on incoming vesicles^22^. In contrast, we discovered that the mammalian exocyst is highly dynamic. Subunit connectivity is conserved between yeast and mammals, but unexpectedly the two subcomplexes, can assemble and localize to the PM independently of each other, and associate with vesicles, yet neither is competent alone to promote vesicle fusion. Moreover, only ~50% of SC1 and SC2 interact in the cytoplasm, and Sec3 is only weakly associated with SC1, demonstrating that mammalian exocyst is not a stable octamer (Fig. 8). Arrival times at the PM are similar for all exocyst subunits and vesicles, consistent with data from HeLa cells^31^ and yeast^21^, but we can detect a small (~80msec) delay between Sec5 (on SC1) and Exo70 (on SC2) suggesting that octameric exocyst complex assembles on the vesicle at or near the PM, perhaps from subcomplexes already attached to the vesicle. The weaker affinity of Sec3 for the complex is consistent with the cryo-EM structure of the yeast exocyst^40^. We propose a comprehensive new model for exocyst interactions (Fig. 8) that while consistent with previous observations expands our view of the dynamics and mechanism of this protein complex.

**Figure 8.**
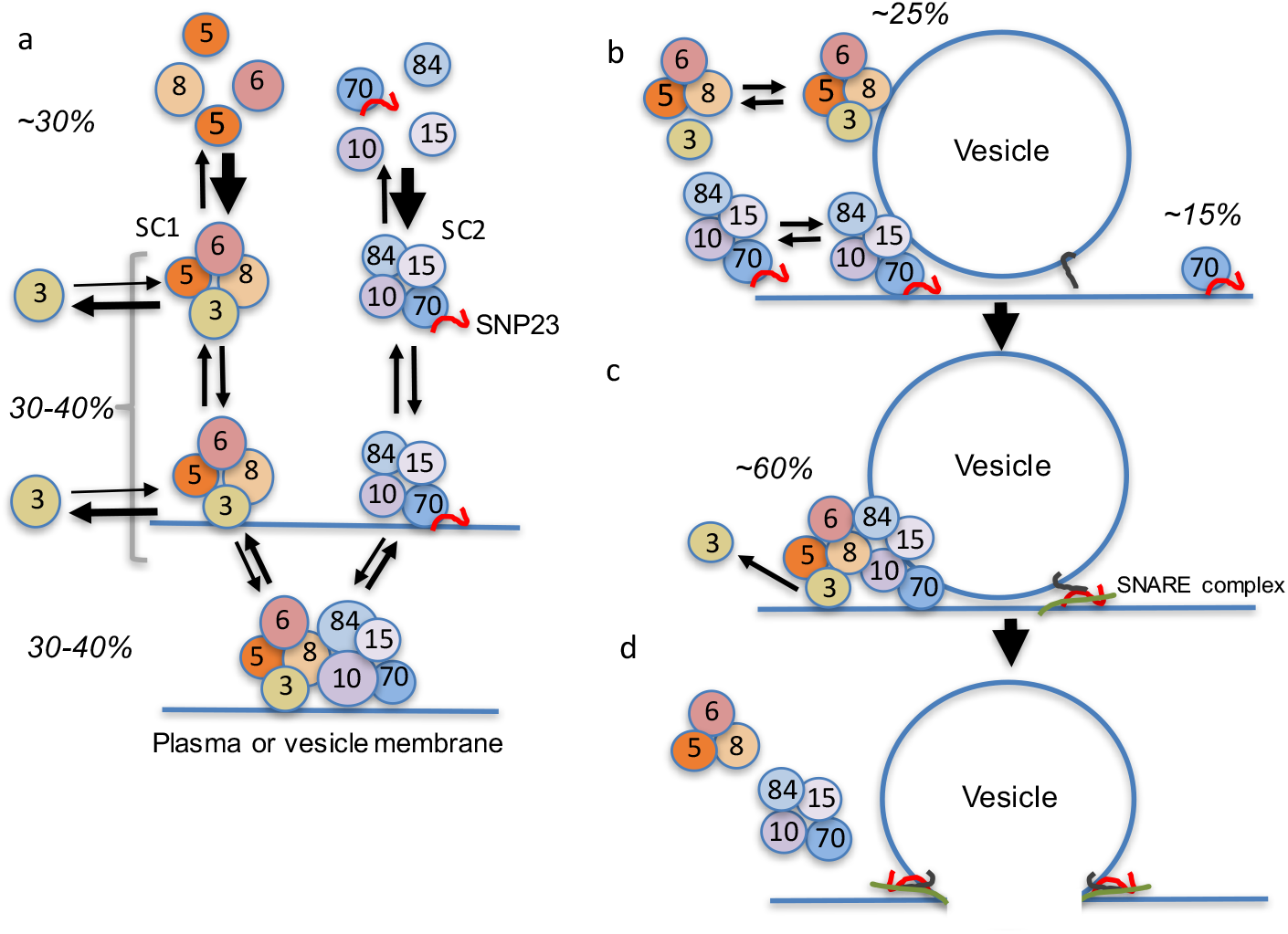
Model for exocyst subunit assembly/disassembly and vesicle tethering. (a) Schematic of subunit interactions within the cell. Subunits are shown as circles, SC1 and SC2 mark the two subcomplexes and arrow weights represent relative on or off rates. SNP23 associates with Exo70 but is not co-precipitated with Sec5. Detailed molecular interactions with membranes are not illustrated and diagram is not to scale. Percentages are the estimated proportions of each state, derived from single molecule counting and FCCS data. Octameric complex abundance = (Sec5+Exo70) colocalization or cross-correlation; tetrameric subcomplex abundance = (Sec5+Sec8) – (Sec5+Exo70); and free subunit abundance = Total – octamer – tetramer. About 50% of exocyst is very slowly diffusing within the cell, likely associated with vesicles. We propose that the individual subcomplexes can interact weakly with membranes but bind membrane more efficiently when assembled into an octamer. Sec3 interacts with SC1 less robustly than do the other subunits. (b) Individual subcomplexes can associate with vesicles at the PM but cannot trigger fusion. Percentages show estimated relative abundances at the membrane, based on TIRFM data. (c) Interaction of the subcomplexes to form octamer stabilizes tethering, and release of Sec3 precedes fusion. (d) Fusion is accompanied by rapid release of the exocyst.

Interestingly, in NMuMG cells we found that the SNARE protein SNAP23 interacts with Exo70 but not with Sec6 or other subunits. No binding of Syntaxins or SM proteins to any of the exocyst subunits was detectable. We speculate that mammalian Exo70 facilitates SNARE complex formation by bringing SNAP23 to the vesicle tethering site, and dissociation of Sec3 from the complex just prior to fusion then enables the SNARE complex to trigger membrane fusion (Fig. 8). However, further studies using endogenously tagged SNARE proteins will be required to test this model.

## Supplemental Information

Supplemental information includes 7 figures and 1 table.

## Acknowledgements

This work was supported by NIH/NIGMS grant (GM070902) to I.G.M., NSERC Discovery Grant (RGPIN 2017 – 06030) to C.C.G., postdoctoral fellowships from the CIHR to S.M.A. and Grant-in-Aid for JSPS Research Fellow (17J40028) to H.N.F. Experiments and data analysis were performed in part through the use of the Vanderbilt Cell Imaging Shared Resources. Mass spectrometry experiments were performed using Vanderbilt MSRC Proteomics Lab. We acknowledge Jorge Rua Fernandez, Rabindra Shivnaraine, Goker Arpag, Marija Zanic, Tomas Kirchhausen and members of the Macara lab for helpful insights. We also thank Yasutsugu Takada, Shigeki Higashiyama and Shinji Fukuda (Ehime University, Japan) for their valuable support.

## Author Contributions

S.M.A., H.N.F., Y.L, and W.H.M. conducted and analyzed the experiments. S.M.A., H.N.F., and I.G.M. designed the experiments. Y.L and C.G performed FCCS data analysis. S.M.A and I.G.M. wrote the paper.

---

**Supplementary Figure 1.**
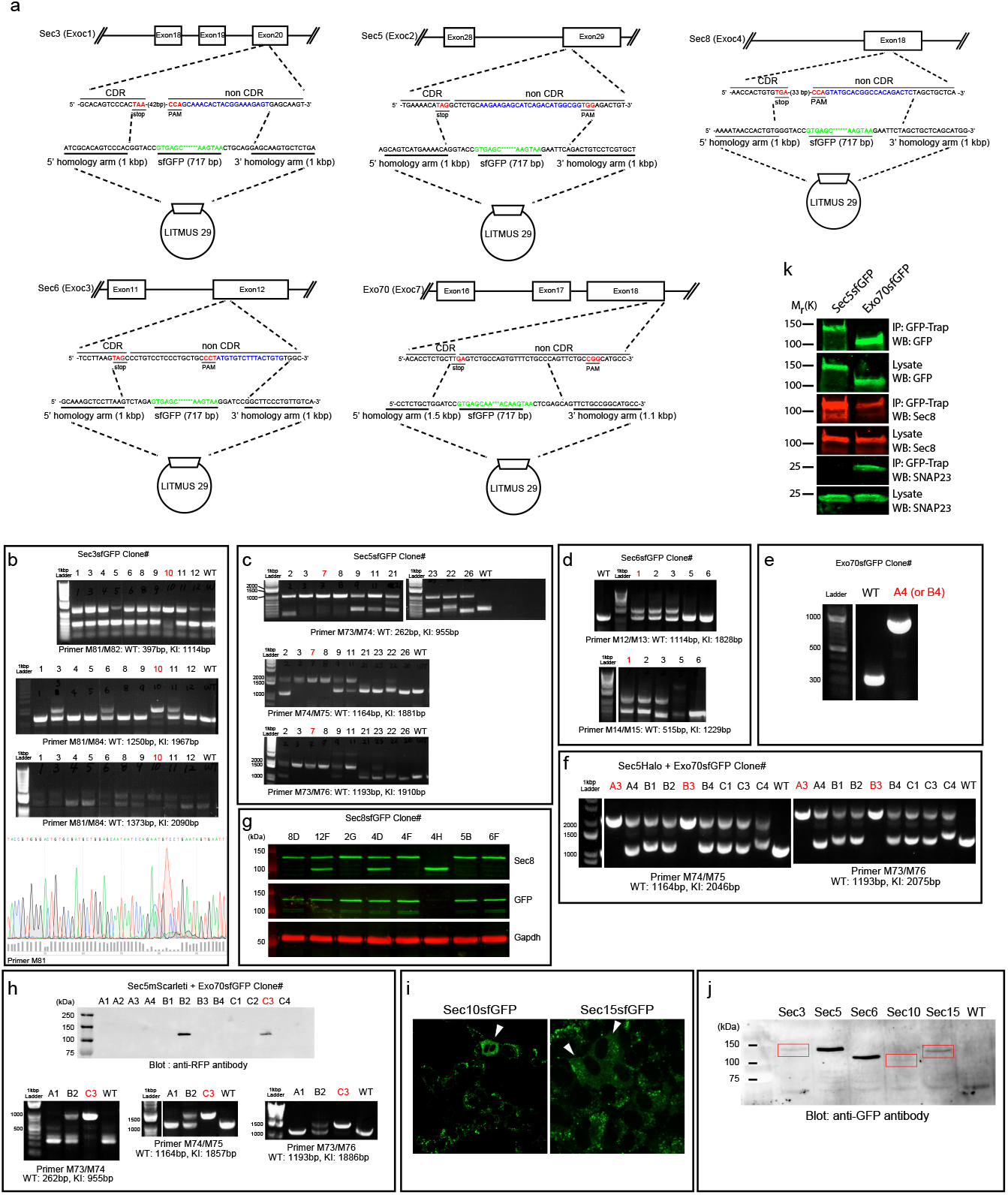
Design and Validation of Gene-Edited Exocyst Cell Lines. (a) Schematic of designs for targeting vectors to insert sfGFP at the C-terminus of exocyst subunits Sec3, Sec5, Sec6 and Exo70 genes. Targeting vector deletes the STOP codon and PAM sequence. (b) Genotypes of Sec3-GFP clones using primer sets listed in STAR methods. Bottom panel shows sequencing of PCR product of the C-terminal region of clone 10. (c-e) Genotypes of Sec5-GFP, Sec6-GFP and Exo70-GFP clones as indicated. (f) Genotype of Sec5-Halo/Exo70-GFP double knock-in clones as indicated. (g) Western blot of Sec8-GFP clones using either anti-Sec8 or anti-GFP antibodies. Gapdh antibodies used to assess loading. (h) Western blot analysis and PCR genotyping of Sec5-mScarlet/Exo70-GFP double knock-in clones. Sec5-mScarlet expression was assessed using anti-RFP antibodies. (i) Confocal images of Sec10-GFP and Sec15-GFP after gene editing and FACS. White arrowheads indicated cells with successful incorporation of sfGFP. (j) Western blot analysis of Sec3-GFP, Sec5-GFP, Sec60GFP, Sec10-GFP and Sec15-GFP cell lines using anti-GFP antibodies. WT = wild type parental cells. (k) Sec5-GFP or Exo70-GFP pulldown using GFP-Trap beads followed by immunoblot analysis to determine binding of native Sec8 (anti-Sec8 antibody) and SNAP23 (anti-SNAP23 antibodies). Pulldowns of Sec5-GFP or Exo70-GFP were assessed using anti-GFP antibodies.

**Supplementary Figure 2.**
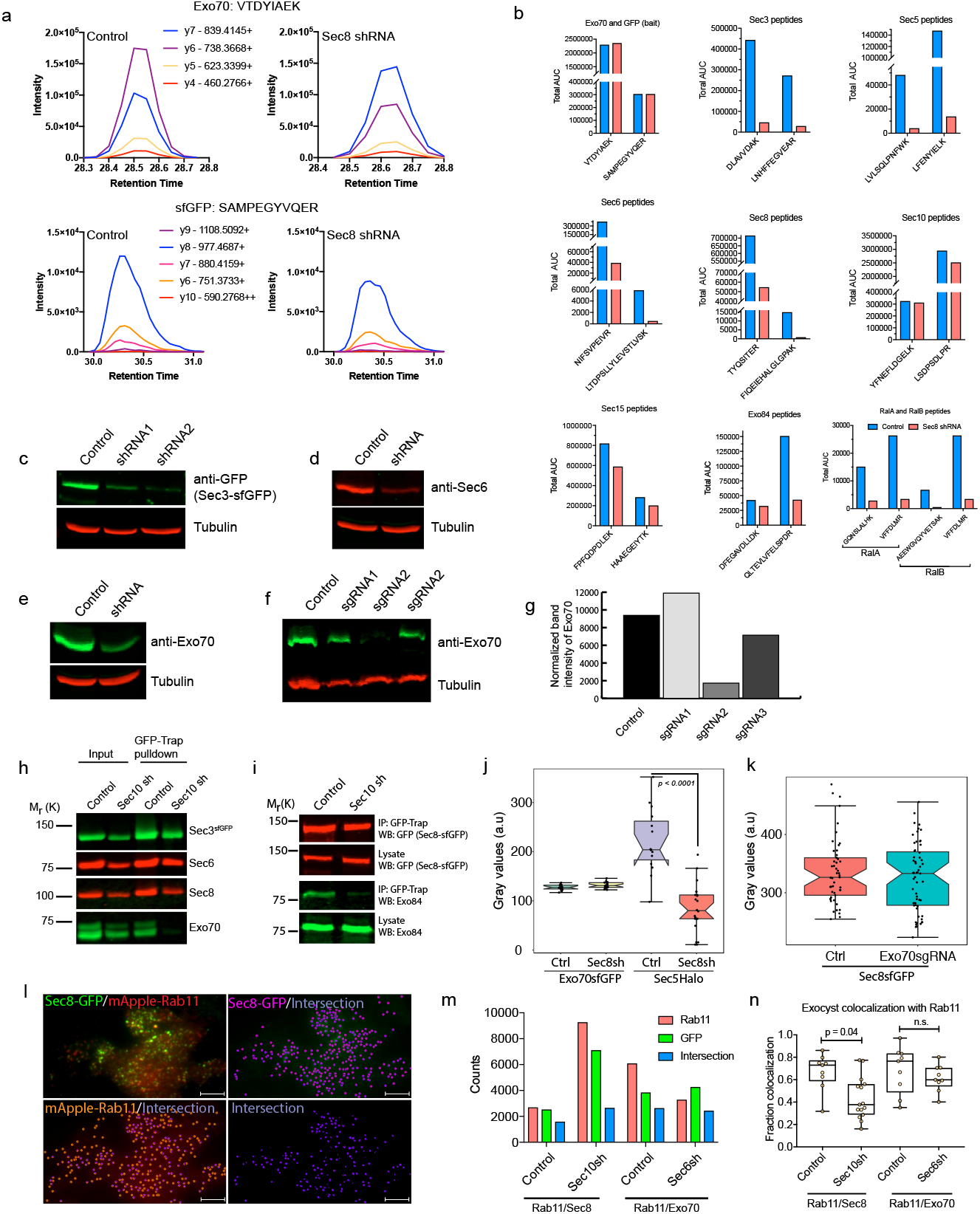
Mammalian Exocyst Subunit Connectivity. (a) Exo70-GFP capture using GFP-Trap nanobodies. Shown are representative MRM spectra of intensity peaks features of the indicated Exo70 and sfGFP peptides in the control or Sec8shRNA treated cells. (b) Quantifications of the area under the curve (AUC) for the indicated peptides for each protein species shown. Control pull-downs are denoted in blue whereas Sec8 depleted conditions are designated in red. (c) Knockdown efficiencies of two independent shRNAs targeting Sec3-GFP. α-Tubulin used as loading control. (d) Knockdown efficiency of Sec6 shRNA. (e) Knockdown efficiency of Exo70 shRNA. (f) Knockout efficiency with three independent guide RNAs targeting Exo70 gene loci in Exon1. (g) Quantification of panel F. (h) GFP-Trap pull-down of endogenous Sec3-GFP from untreated or Sec10 shRNA treated NMuMG cells. Western blots were immunoblotted with anti-GFP to detect Sec3-GFP, or with anti-Sec6, anti-Sec8 and anti-Exo70 antibodies to assess co-precipitations of unlabeled exocyst subunits. (i) GFP-Trap pull-down of endogenous Sec8-GFP from untreated or Sec10 shRNA treated NMuMG cells. Blots were probed with anti-GFP to detect Sec8-GFP, or with anti-Exo84 antibodies to assess amount of co-precipitation of endogenous unlabeled Exo84. (j) Quantification of fluorescence intensities from TIRFM images of Sec5-Halo and Exo70-GFP double knock-in cells treated with Control or Sec8 shRNAs. Sec5-Halo was labeled with JF585 Halo ligand. (k) Fluorescence intensity quantifications of Sec8-GFP from TIRFM images in untreated or Exo70 knockout cells. (l) Example of Sec8-GFP and mApple-Rab11 coincidence. Cells were transduced with mApple-Rab11 lentivirus. Spot diameters in the range of 0.30-0.35μm was used to identify GFP and Rab11 and the fraction of the particle that are both red and green were determined using NIS Elements spot detection algorithm. Scale bar = 5μm. (m) Total number of particles analyzed for Rab11, exocyst subunits and fraction that co-localize. (n) Quantification of the fraction of exocyst subunits Sec8-GFP or Exo70-GFP that coincide with mApple-Rab11. Ordinate axis = intersection/GFP. Cells were treated with scrambled, Sec10 or Sec6 targeting hairpins. Center lines show the medians; box limits indicate the 25th and 75th percentiles as determined by R software; whiskers extend 1.5 times the interquartile range from the 25th and 75th percentiles, data points are plotted as dots. Statistical significance was assessed using one-way ANOVA followed by Sheffe’s multiple comparison tests.

**Supplementary Figure 3.**
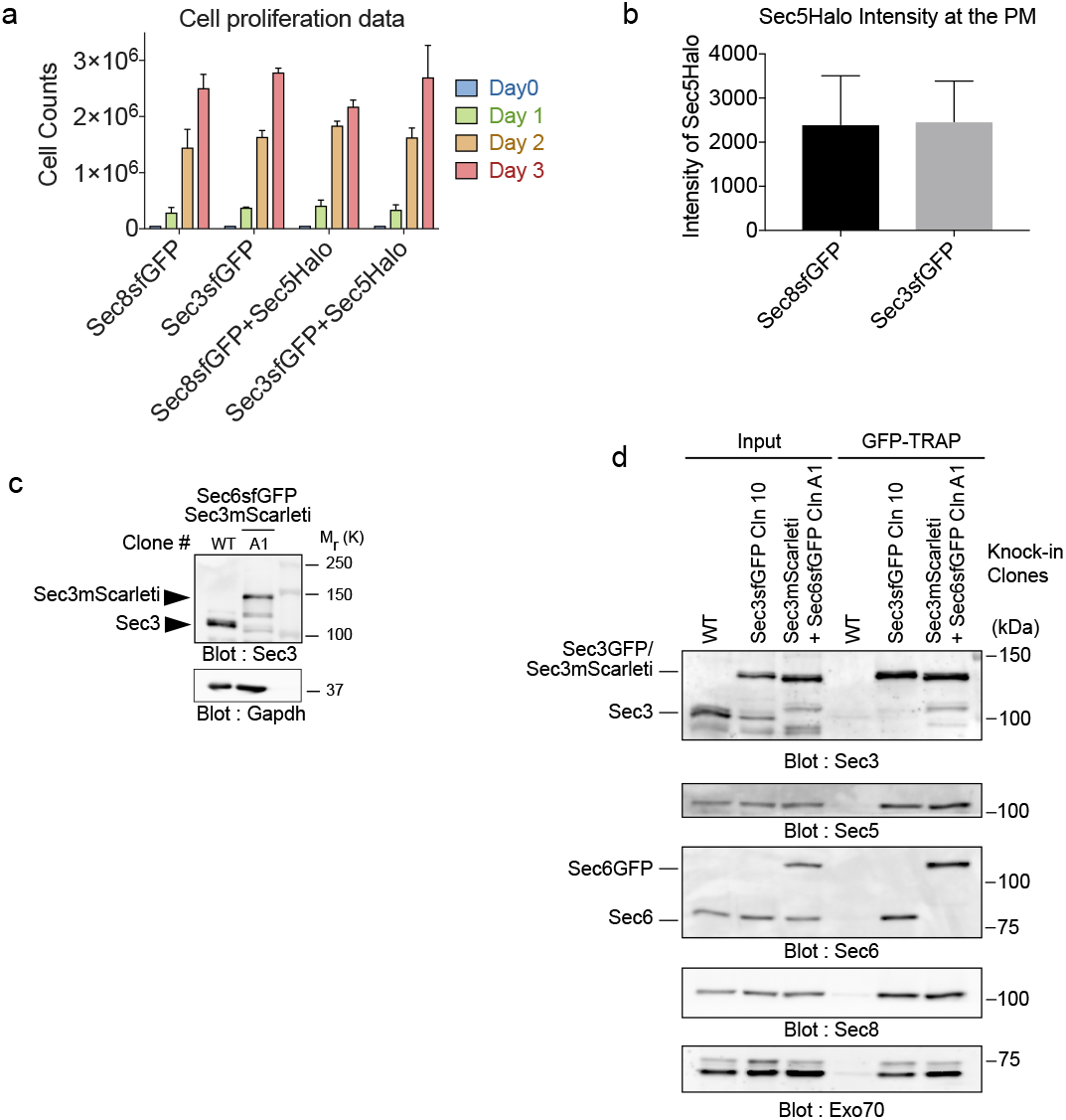
Functionality of Exocyst Knock-in Cell-Lines. (a) Cell proliferation of Sec8-GFP, Sec3-GFP, Sec8-GFP + Sec5-Halo and Sec3-GFP + Sec5-Halo knock-in cells over three days. Error bars indicate ± *s.d.* Statistical analysis was using one-way ANOVA to compare groups for each day. (b) Comparison of Sec5-Halo fluorescence intensity at the bottom PM (PM) in double knock-in cells. Intensities were measured using TIRFM. Halo tag was labeled with JF585-HTL. (c) Western blot analysis of Sec3-Scarlet+ Sec6-GFP double knock-in cells. Clone A1 is homozygous for Sec3. WT = wild-type parental NMuMG cells. (d) GFP-Trap pull-down and western blots analysis from double knock-in cell lines expressing Sec3-GFP, or Sec3-Scarlet + Sec6-GFP. WT = wildtype cells, Cln = clone.

**Supplementary Figure 4.**
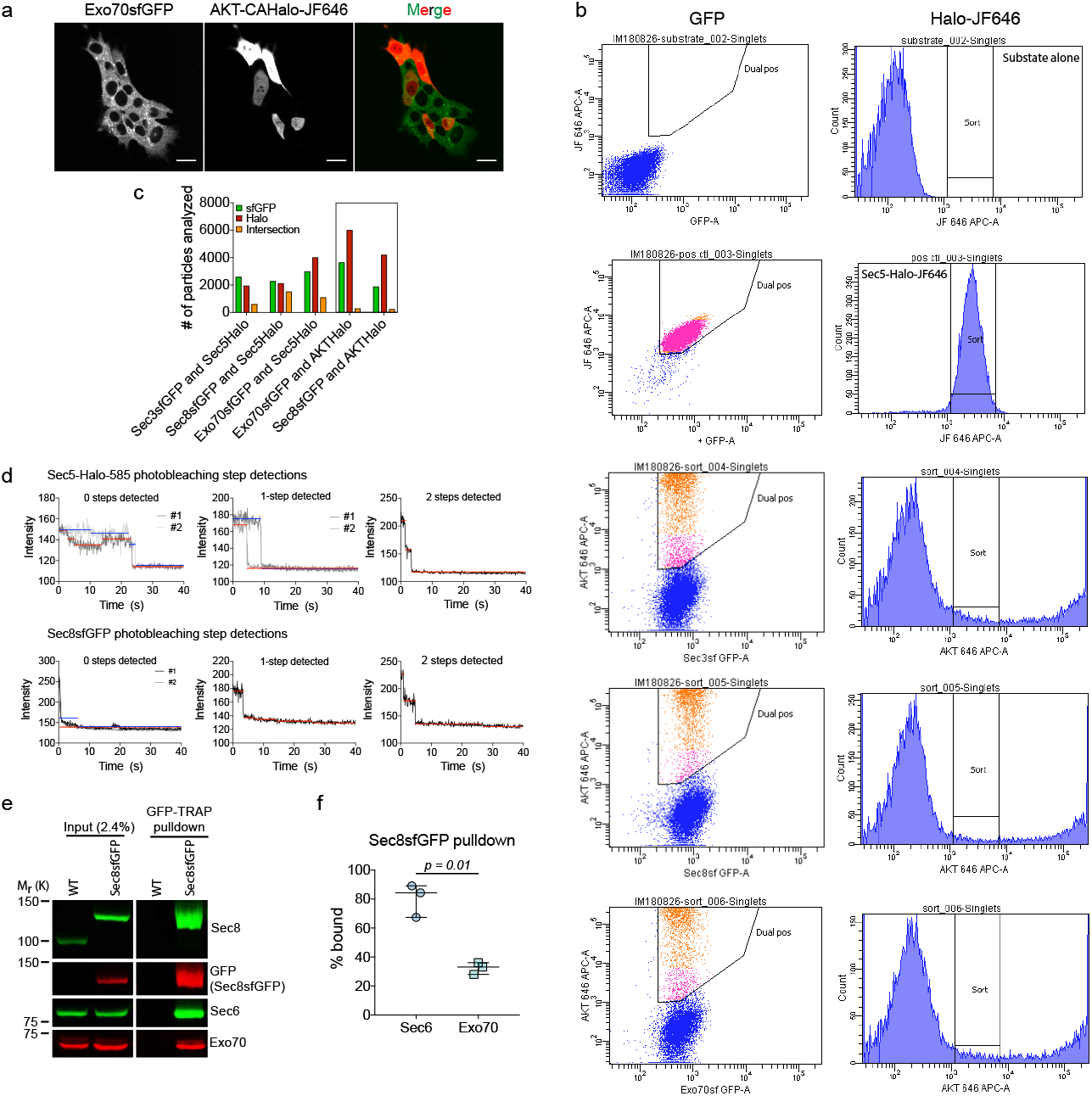
Fractional Interactions of Cellular Pools of Exocyst subunits. (a) AKT-Halo was expressed in Exo70-GFP cells as a negative control for interactions. Halotag was labeled with JF-646 HTL and and imaged to look at the distribution of expression prior to cell sorting. Scale bar = 20μm. (b) Sec3-GFP, Sec8-GFP and Exo70-GFP knock-in cells were transduced with lentivirus expressing AKT-Halo, and sorted for similar expressed levels than Sec5-Halo. Halo was labeled with JF-646 HTL. (c) Graph shows number of molecules analyzed in the data shown in Figure 5d. (d) Representative images of 0, 1 or 2 photobleaching steps detected by the step detection algorithm. (e) Representative experiment where Sec8-GFP was captured using GFP nanobodies and immunoblotted with anti-Sec8, anti-GFP, anti-Sec6 or anti-Exo70 antibodies. Blots on the left show 2.4% input of the total lysate and the blot of the right shows pulldown with nanobodies. (f) Quantification of the percentage of Sec6 and Exo70 bound to Sec8 from 3 independent experiment experiments.

**Supplementary Figure 5.**
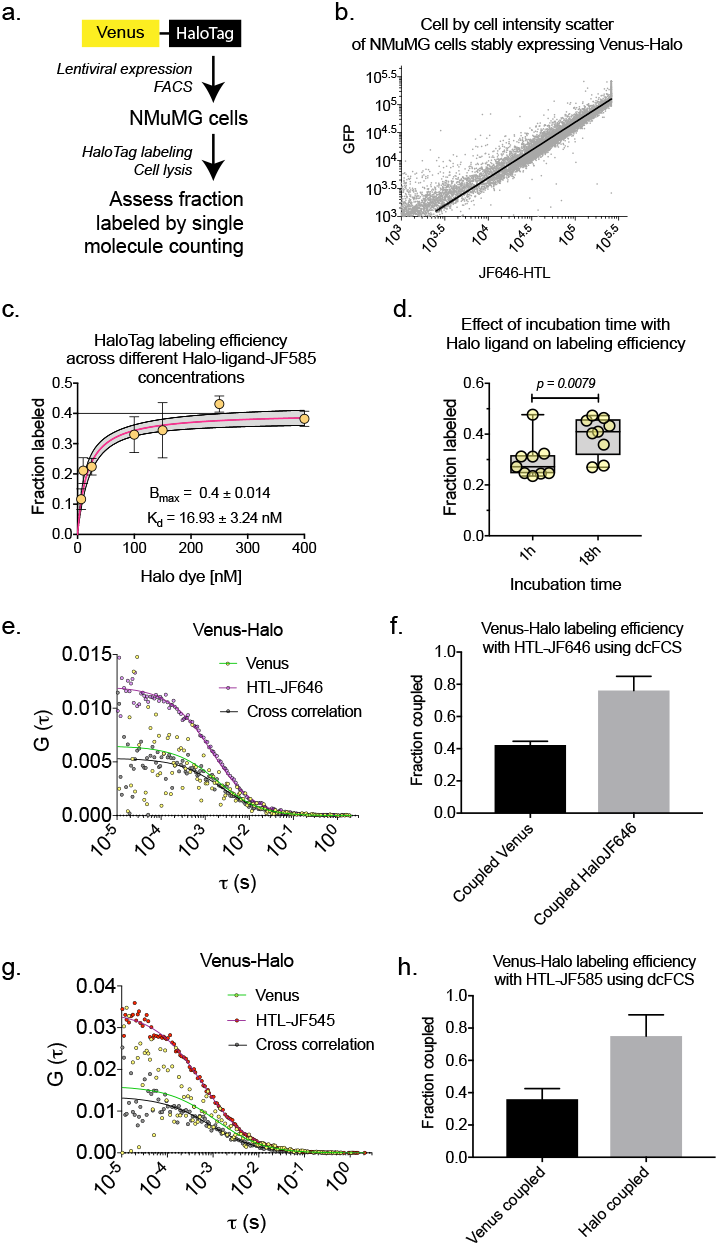
HaloTag Labeling Efficiency. (a) Schematic showing experimental design to determine Halo labeling efficiency in NMuMG cells. Halo fused to the C-terminus of of YFP was expressed in NMuMG cells using lentiviral expression system, and subsequently sorted for positive cells. (b) Correlation between fluorescence intensities of YFP and Halo labeled with JF-646-HTL. (c) Halo labeling efficiency across different concentrations (6.25, 25, 100, 150, and 400 nM) of HaloTag ligand conjugated to JF585 dye in NMuMG cells for 1.5h. (d) Halo labeling efficiency after NMuMG cells were labeled with 100nM of HaloTag ligand conjugated to JF585 for 1h or 18h. (e) Dual-color FCCS measurement of Venus-Halo labeled with 150nM HTL-JF646 for 2h in NMuMG cells. (f) Quantification of the fraction of Venus-Halo that are labeled with HTL-JF646. (g) Dual-color FCCS measurement of Venus-Halo labeled with 150nM HTL-JF585 for 2h in NMuMG cells. (h) Quantification of the fraction of Venus-Halo that are labeled with HTL-JF585.

**Supplementary Figure 6.**
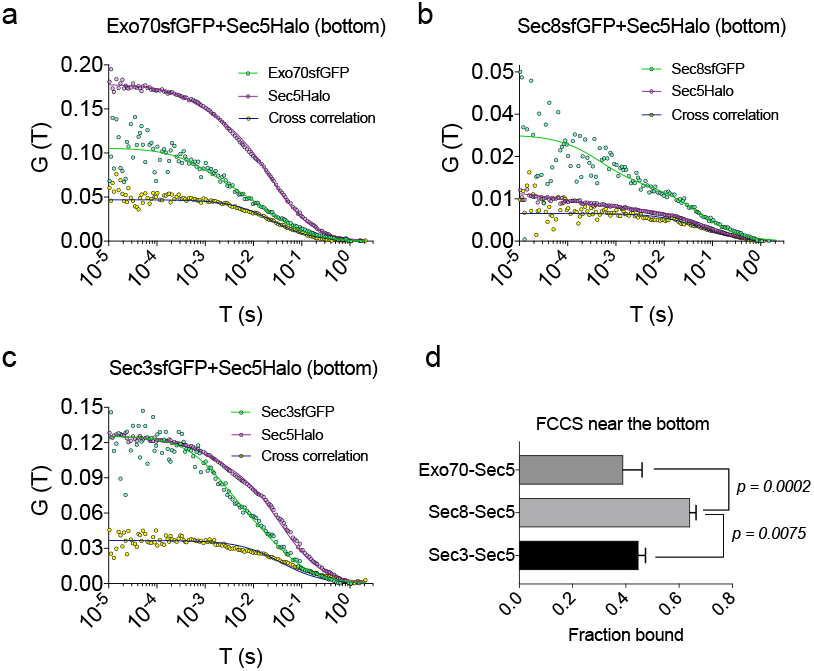
Dual-Color FCCS Measurements of Exocyst Subunits near the base of the cells. (a) Sec5-Halo and Exo70-GFP, (b) Sec5-Halo and Sec8-GFP and (c) Sec5-Halo and Sec3-GFP FCS measurements near the base of the cells. (d) Statistics of fraction of GFP-tagged exocyst subunits near the bottom membrane of the cells. Sec8-GFP+Sec5-Halo: 64% ± 2.2%, Sec3-GFP+Sec5-Halo: 44% ± 2.6%, Exo70-GFP+Sec5-Halo: 39% ± 7.0% (mean ± *s.e.m*.). These numbers are probably underestimates as exocysts on the membrane are less diffusive. Also the measurements likely include both plasma membrane bound fraction as well as cytoplasmic fraction. HaloTag was labeled using JF646 Halo ligand (200 nM for 1.5h).

**Supplementary Figure 7.**
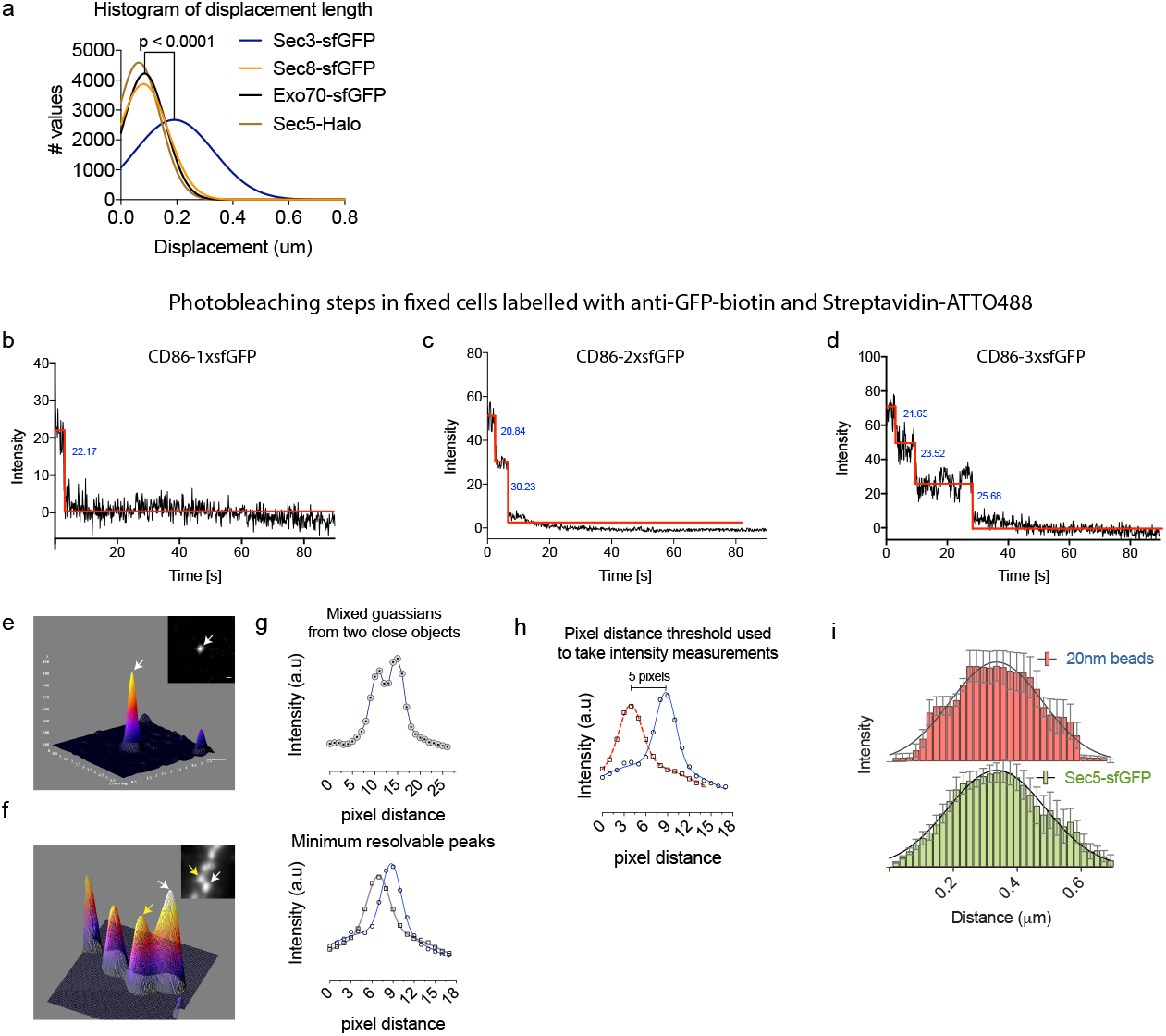
Determination of Criteria Used to Count Exocyst Subunit Molecules from Fluorescence Intensities. (a) Distribution of vector displacements between the first and last frames of the particle trace. Data are shown from >20,000 object for each exocyst subunit as indicated. Statistical significance was measured by Kruskal-Wallis non-parametric test followed by Dunn’s post-hoc test. Photobleaching steps of (B) CD86-1xsfGFP (C) CD86-2xGFP and (D) CD86-3xsfGFP in cells fixed with 3.7% paraformaldehyde followed by methanol and immunolabeled with anti-GFP-biotin antibodies and Streptavidin-ATTO488 dye. (e) Fluorescence landscape showing point-spread function of a single tetraspec bead spread on glass. (f) Fluorescence landscape of multiple tetraspec beads close together. Arrows point to two close but resolvable peaks from which individual intensities could be measured. (g) 2D intensity plots of the peaks described in F. (h) Intensity trace shown in panel is shifted to show minimal separation in pixel distance (5 pixels) that was used to measure intensities of the objects in cells. (i) Distributions of average intensity traces over a 20nm bead (n=3) or a Sec5-GFP particle at vesicle fusion sites (n=5).

## Methods

### Plasmids constructs and other reagents

pIRESpuro2-CD86-mEos2 and pIRESpuro2-CTLA4-mEos2 were gifts from Mike Heilemann (Addgene plasmids # 98284 and 98285). mEos2 was replaced with superfolder GFP coding sequence (sfGFP), which was PCR amplified from sfGFP-N1, a gift from Micheal Davidson (Addgene plasmid # 54737), with flanking XmaI/NotI restriction enzyme sites and inserted at the AgeI/NotI site of the plasmid in frame with CD86. As such, the AgeI site at the beginning of sfGFP was destroyed and a new AgeI site created at the end of sfGFP before the stop codon, to enable insertion of subsequent tandem sfGFP coding sequences in frame. pEGFP-VAMP2 was a gift from Thierry Galli (Addgene plasmid # 42308). TfR-pHuji was a gift from David Perrais (Addgene plasmid # 61505). Superecliptic pHluorin (pHluorin2) was PCR amplified from VV063: 1xCox8-superecliptic pHluorin in fck, a gift from Adam Cohen (Addgene plasmid # 58500) to make VAMP2-pHluorin2. Halo-tag coding sequence was synthesized as a geneblock (IDT) to make VAMP2-Halo. AKT-Halo construct was generated by replacing mApple with HaloTag using the EcoRI/NdeI restriction sites at the C-terminal end of a pLVTHM-AKT-CA construct described previously ^30^. For CRISPR/Cas9-mediated gene editing, locus-specific 5’ and 3’ homology arms were synthesized as gBlocks double-stranded gene fragments (IDT), or amplified by PCR. These gene fragments were designed to remove the stop codon from the gene ORF, delete the PAM sequence recognized by the cognate sgRNA, and to attach an in-frame tag (sfGFP, HaloTag or mScarleti). They were then cloned into LITMUS29 vector together with the appropriate tag either using Gibson assembly (NEB) or by conventional cloning using the following restriction sites: KpnI/PstI (Sec3), KpnI/EcoRI (Sec5), XbaI/BamHI (Sec6), KpnI/EcoRI (Sec8), BamHI/XhoI (Exo70).

The following hairpin RNA clones were purchased from the Sigma MISSION shRNA library: mouse Sec3 shRNA clones TRCN0000254147, TRCN0000254146 and TRCN0000254145, Sec6 shRNA clones TRCN0000111629, mouse Sec8 shRNA clones TRCN0000307390 and TRCN0000298307, mouse Sec10 shRNA clone TRCN0000093547, and Exo70 shRNA clone TRCN0000376796. Silencing effiiciencies of the shRNA were verified by immunoblotting analysis compared to a control shRNA targeting Luciferase. Exo70 sgRNA (5’-CACCGGTTGTCTGGCAGCTGGCTA-3’) was cloned into lentiCRISPR-v2 at the BsmBI restriction site.

For immunoblotting, we used the following antibodies: mouse anti-Sec8 (Clone 14/Sec8, BD Biosciences), mouse anti-Sec6 (Clone 9H5, Novus Biologicals), Rabbit anti-Exo70 (Bethyl Laboratories), Rabbit anti-GFP (ThermoFisher Scientific), chicken anti-GFP (Abcam), goat anti-GFP-biotin (Abcam), rabbit anti-RFP (Rockland), mouse anti-α-tubulin (Clone DM1A, Sigma-Aldrich), rabbit anti-GAPDH (Clone 14C10, Cell Signaling Technology), mouse anti-RalA (Clone 8/Ral A, BD Transduction Laboratories). GFP-Trap^®^_MA beads were purchased from ChromoTek GmbH.

Other materials used in this paper were as follows. Q-VD-Oph (Sigma-Aldrich); ProLong^®^ Live Antifade Reagent (Thermo Fisher Scientific). Trolox (Acros Organics) was used for live cell imaging. FluoroBright DMEM medium for live cell imaging was from Life Technologies. Fetal Bovine Serum (FBS) was from Atlantic Biologicals. JF585-Halo and JF646-Halo dyes were gifts from Luke Lavis, Janelia Farms HHMI Institute^47^.

### Cell culture

NMuMG (ATCC CRL-1636) and HEK293T (ATCC CRL-3216) cells were obtained from ATCC. Cells were cultured in Dulbecco’s Modified Eagle Medium (Life Technologies), supplemented with 10% FBS and 1X Penicillin/Streptomycin (Life Technologies) and maintained in culture as suggested by ATCC.

### Lentiviral transductions and transient transfections

Lentivirus was produced by transfecting HEK293T cells with lentiviral packaging vectors pMD2.G and psPAX2 using calcium phosphate precipitation. Lentiviral transductions were with virus-conditioned medium collected from HEK293T cells 48h post-transfection. Xfect transfection reagent (Takara) was used to transiently transfect NMuMG cells, to create exocyst knock-in cell lines.

### CRISPR/Cas9-mediated generation of knock-in cell lines

NMuMG cells were plated at 10^5^ cells/well in 6 well plate 24 h before transfection. Cells were transfected with targeting vector (LITMUS29 backbone), sgRNA (Addgene plasmid #41824) and Cas9 expression vector (pCMVsp6-nls-hCas9-nls) a gift from Li-En Jao (UC – Davis), at an equimolar (550 fmol) ratio using Xfect transfection reagent according to the manufacture’s protocol. The GFP positive cells were subsequently sorted on a FACSAriaIII (BD) 5 d post-transfection. After single cell cloning, insertions were confirmed by immunoblot analysis and PCR-based genotyping.

### Stable cells

NMuMG cells transduced with shRNAs or sgRNA were selected using puromycin (2μg/ml). Cells infected with Vamp2-pHluorin, Vamp2-Halo or AKT-Halo were selected using Blasticidin (10μg/ml) or FACS.

### Immunoprecipitation and quantitative western blot

NMuMG CRISPR cells lines expressing sfGFP-tagged exocyst subunits were grown to confluency in 100mm cell culture dishes. Cells were washed with PBS and lysed in 1ml lysis buffer containing 20mM HEPES; pH 7.4, 50mM NaCl, 2mM EDTA, 0.1% Triton X-100, supplemented with cOmplete^™^ mini EDTA-free protease inhibitor cocktails (Roche) and PhosStop (Roche). Cells were lysed in a rotator for 15 min at 4°C and debris removed by centrifugation at 16,000xg. For quantitative immunoprecipitation protein amounts were measured using a BCA assay (ThermoFisher Scientific) and small amount of lysates from each step (input, unbound, wash) were retained for analysis. Affinity purifications were performed with GFP-Trap_MA beads. Quantitative immunoblot analysis used secondary antibodies conjugated to IRDye^^®^^ 680RD or 800CW (LI-COR), imaging blots with Odyssey CLx Infrared Scanner (LI-COR).

### Affinity purification for mass spectrometry

Gene edited NMuMG cell lines were grown to confluency in 4 to 5 150mm cell culture dishes. Cells were washed 3x with 20ml PBS and lysed in 10ml lysis buffer described above, supplemented with cOmplete^™^ mini EDTA-free protease inhibitor cocktails (Roche), PhosStop (Roche) and NaF (100mM; Sigma-Aldrich). Cells were harvested using a cell scraper followed by immediate freezing in liq N_2_. Lysates were thawed and briefly sonicated on ice using a microtip sonicator (20s; Vibra-Cell), followed by centrifugation at 16,000xg for 10 min at 4°C to remove debris. Supernatants were transferred into 15ml conical tubes and baits captured on GFP-Trap^^®^^_MA beads by end over end mixing at 4°C for 30 min. Beads were separated using Magnetic Particle Concentrator (ThermoFisher Scientific), unbound supernatant aspirated followed by 3 x 1.5ml washes in the lysis buffer. Samples were eluted in 2x Laemmli sample buffer and heated to 95°C for 15 min and subsequently processed for mass spectrometry (MS) analysis).

### Sample preparation for multiple reaction monitoring mass spectrometry

Affinity purification was performed as mentioned above and relative abundance of each exocyst subunits binding to the bait was compared across label free samples by multiple reaction monitoring mass spectrometry (MRM-MS). Quantified stable-isotope-peptides for quantitative MRM were obtained from JPT Peptide Technologies GmbH.

The quantified stable-isotope-labeled peptide standards were spiked into the sample of interest and peptides from the endogenous exocyst subunits quantified from the ratio of the labeled peak heights to the unlabeled.

### Mass spectrometry

*LC-MS/MS:* Exocyst protein immunoprecipitations were run on a NuPAGE Bis-Tris gel, were stained with Novex colloidal Coomassie stain (Invitrogen), and destained in water. Coomassie stained gel regions were cut from the gel and diced into 1mm^3^ cubes. Proteins were treated for 30 minutes with 45 mM DTT, and available Cys residues were carbamidomethylated with 100mM iodoacetamide for 45 min. Gel pieces were further destained with 50% MeCN in 25mM ammonium bicarbonate, and proteins were digested with trypsin (10ng/uL) in 25mM ammonium bicarbonate overnight at 37°C. Peptides were extracted by gel dehydration with 60% MeCN, 0.1% TFA, the extracts were dried by speed vac centrifugation, and reconstituted in 0.1% formic acid.

Peptides were analyzed by LC-coupled tandem mass spectrometry (LC-MS/MS). An analytical column was packed with 20cm of C18 reverse phase material (Jupiter, 3 μm beads, 300Å, Phenomenox) directly into a laser-pulled emitter tip. Peptides were loaded on the capillary reverse phase analytical column (360 μm O.D. x 100 μm I.D.) using a Dionex Ultimate 3000 nanoLC and autosampler. The mobile phase solvents consisted of 0.1% formic acid, 99.9% water (solvent A) and 0.1% formic acid, 99.9% acetonitrile (solvent B). Peptides were gradient-eluted at a flow rate of 350 nL/min, using a 110-minute gradient. The gradient consisted of the following: 1-3min, 2% B (sample loading from autosampler); 3-88 min, 2-40% B; 88-98 min, 40-95% B; 98-99 min, 95% B; 99-100 min, 95-2% B; 100-110 min (column re-equilibration), 2% B. A Q Exactive Plus mass spectrometer (Thermo Scientific), equipped with a nanoelectrospray ionization source, was used to mass analyze the eluting peptides using a data-dependent method. The instrument method consisted of MS1 using an MS AGC target value of 3e6, followed by up to 20 MS/MS scans of the most abundant ions detected in the preceding MS scan. A maximum MS/MS ion time of 60 ms was used with a MS2 AGC target of 1e5. Dynamic exclusion was set to 15s, HCD collision energy was set to 28 nce, and peptide match and isotope exclusion were enabled. For identification of peptides, tandem mass spectra were searched with Sequest (Thermo Fisher Scientific) against a *Mus musculus* database created from the UniprotKB protein database (www.uniprot.org). Variable modification of +15.9949 on Met (oxidation) and +57.0214 on Cys (carbamidomethylation) were included for database searching. Search results were assembled using Scaffold 4.3.2. (Proteome Software).

### MRM-MS

Peptides for each protein were selected based on their appearance in data dependent analyses and then optimized for the most useful transitions to monitor. Heavy-labeled peptide internal standards were synthesized by jpt (SpikeTides TQL peptides, jpt, Berlin, Germany) which contained isotopically-labeled terminal arginine or lysine residues (^13^C and ^15^N) and a trypsin-removable C-terminal tag. These isotopically-labeled peptides were digested separately and then spiked into samples at approximately endogenous levels after in-gel digestion of sample proteins. Skyline software (University of Washington, MacCoss lab) was used to set up scheduled, targeted MRM methods monitoring four to five MRM transitions per peptide. A final MRM instrument method including the isotopically labeled standards and encompassing a 9-minute window around the retention time of each peptides was performed using a 40 mm by 0.1 mm (Jupiter 5 micron, 300A) kasil fritted trap followed by a 250 mm by 0.1 mm (Jupiter 3 micron, 300A), self-packed analytical column coupled directly to an TSQ-Vantage (ThermoFisher) via a nanoelectrospray source. Peptides were resolved using an aqueous to organic gradient flowing at 400 nl/min. Q1 peak width resolution was set to 0.7, collision gas pressure was 1 mTorr, and utilized an EZmethod cycle time of 3 seconds.

### Single molecule approach to assess protein-protein interactions

#### Surface cleaning

No. 1.5 glass coverslips (Electron Microscopy Sciences) and quartz slides were (Ted Pella Inc.) rinsed in 100% ethanol, followed by acetone and sonicated for 15 min. Coverslips and slides were then put in 1M KOH (Sigma-Aldrich) and sonicated for an additional 30 min, followed with 3 rinsed in ultrapure water, and additional rinses in acetone. The materials were further cleaned in 0.5 vol-% Helmanex III solution for 30 minutes followed by rinsing in ultrapure water three times. Surfaces were then sonicated for 10 min in 99.9% HPLC-grade methanol (Fisher Chemical). The surfaces were finally torched by swiping three times over a Bunsen burner.

#### Labeling, cell lysis and imaging

Approximately 300,000 cells from 24-well plates were used. Cells were first labeled with JF585-Halo dye for 90 min, followed by extensive 5x washes with PBS to remove any unbound dye. Cells were then lysed in ice-cold lysis buffer (20mM Hepes, 50mM NaCl, 2mM EDTA, 0.15% Triton X-100), centrifuged at 13,000xg for 5 min to remove debris and a fraction of the cleared lysate was spread in between clean quartz slides and a glass coverslip for imaging. Images were taken using a Nikon TIRF microscope, Apo-TIRF 100x/1.49 oil immersion lens online with a Photometrics Prime 95B sCMOS camera. Both GFP and JF585-Halo were simulatenously excited using AOTF driven Nikon LUN-F multi-excitation diode laser lines firing 488nm and 561nm lasers and images collected on the camera using an Optosplit III setup (Cairn Research). Images were taken at 12.5Hz for 40 sec. Photobleaching steps were analyzed in Matlab using the algorithm described previously (Shivnaraine et al., 2016). Fraction bound was determined according to colocalization of green and red particles, analyzed in FIJI.

#### Determining HaloTag labeling efficiency

To determine the efficiency of HaloTag labeling in NMuMG cells we created a Venus-HaloTag fusion construct in lentiviral vector pLV-Venus. HaloTag coding sequence was inserted C-terminally to Venus between restriction sites BamHI/NdeI. Venus-Halo was expressed in NMuMG cells to test HaloTag labeling efficiency on incubation for 1.5h with different concentration of JF585-labeled-HaloTag ligand, or incubating cells with a single concentration of the ligands for 1h or 18h. Labeling efficiency was determined by spreading the cleared cell lysates as described above and calculated according to the expression:

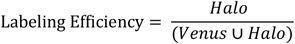

### Live cell TIRFM imaging and analysis

TIRF images were collected with an Andor Zyla sCMOS on Nikon Ti-E or a Photometrics Prime 95B mounted on a Nikon Ti-2 microscope, using an Apo TIRF 60x/1.49 oil immersion lens. The Nikon Ti-E TIRF microscope was equipped with multi-excitation diode laser lines (405nm,488nm, 561nm and 647nm) with triggering capabilities for fast dual color acquisitions, as described above. Image acquisitions were performed using NIS Elements Advanced Research imaging software [version 4.60.00 (Build 1171) Patch 02]. Images were processed and analyzed with Elements or FIJI.

For analysis of single fusion events, each acquired image sequence was manually reviewed multiple times to visually identify vesicle fusion events as demarcated by Vamp2-pHluorin or TfR-pHuji fluorescence intensity flashes. Coordinates of the vesicle fusion events were marked by a circular ROI. GFP or mApple particle durations were analyzed using NIS-Elements. Circular ROIs ~1 μm were drawn inside the initial ROIs and were used to calculate the intensities of sfGFP and pHuji. Arrival and departure of sfGFP were determined based on their increase and decrease in intensities, respectively. Kymographs of sfGFP and pHuji were generated and analyzed using NIS-Elements.

Particle coincidences was measured using NIS-Element’s Binary/spot detection algorithm. Spot diameter parameters were set between 0.27-0.35μm, and the contrast was adjusted to detect objects above background for these analysis. The number of spots that were coincidence was measurement subsequently.

Photobleaching step detection in Figure 4 and Figure S4, was done in Matlab using an algorithm described previously ^48^. Exocyst molecule numbers were estimated from GFP intensity standards. One, 2 or 3 sfGFP were fused to the C-terminus of monomeric CD86 and expressed in HEK293T cells ^49^. Intensities were measured at a fixed optimal camera and laser setting. Intensities of the molecules were matched to photobleaching steps and fitted to a linear regression model. The same instruments settings and the region of the image with uniform intensities were then used to measure exocyst molecule intensities during vesicle fusions. The peak intensities of single resolved sfGFP-exocyst particles at vesicle fusion sites, indicated by TfR-pHuji, were then measured and the number of molecules/particle determined from the regression equation. Supplementary Fig. 6e-i further explains the criterions used for fluorescence intensity measurements.

Mean squared displacements of the different exocyst subunits from TIRFM images were measured using Imaris image analysis software (Bitplane) tracking algorithm. For two-color tracking, each channel was individually processed and overlaid to assess differences in overall diffusion patterns.

### Dual Color Fluorescence Cross-Correlation Spectroscopy in Live Cells

#### Experimental setup

dcFCS measurements were performed on a custom-built instrument using a hardware correlator ^50^. The system is a confocal microscope built with multi-color excitation and detection and with real-time multi-channel correlation analysis capabilities. sfGFP and JF646 were excited simultaneously by a blue laser (TECBL-488nm, WorldStarTech) and a red laser (TECRL-633nm, WorldStarTech). Excitation power at the sample was set in the range of 0.1 – 5 *μ*W, corresponding to an average intensity of 0.05 – 2.5 kW/cm^2^ in the diffraction-limited confocal area. Fluorescence signals from sfGFP and JF646 were separated by a dichroic filter (FF585-Di01, Semrock) and directed through a green bandpass filter (FF01-512/25, Semrock) in the case of eGFP or a red longpass filter (LP02-647RS, Semrock) in the case of JF646. Each signal was split using a 50/50 non-polarizing beamsplitter. The four signals were collected using four avalanche photodiode (APD) detectors (SPCM-AQR-13-FC, PerkinElmer Inc.) and processed by a multichannel hardware correlator (Flex11-8Ch, correlator.com). The correlator outputs up to four different intensity correlation curves between any two detection channels. For typical live cell measurements, at least 10^7^ photons were collected from each cell to construct the FCS curves.

#### Live cell measurements

Experiments on NMuMG cells were performed using glass-bottomed Petri-dishes (MatTek), in which the cells grow on a round coverslip located in the center at the bottom of the dish. All imaging was performed in FluoroBrite DMEM media (Thermo Fisher Scientific). Confocal scan imaging was employed to search for cells that exhibit an optimal fluorescence signal of 5 - 100 kHz at a laser excitation intensity of 50 - 250 W/cm^2^. The software-controlled raster scan was performed using a piezo nano-positioning stage (Nano-LP100, Mad City Labs Inc.), which has a 100-μm scanning range on each of the three axes. There typically was less than one cell suitable for FCS measurements within the 100-μm field of view. A mechanical translation stage was used to bring new 100-μm regions of the dish into the field of view of the microscope. This allowed for the survey of multiple cells in multiple regions within a single dish, thus improving efficiency of data collection and the statistical reliability of data analysis.

### Model-fitting for FCS

Correlation curves from dcFCS were analyzed in MATLAB using a custom-written program based on the Marquardt-Levenburg algorithm. The data were weighted according to the standard deviation at each point, as determined from multiple repeated readings. Rhodamine 110 and Alexa Fluor 647 were measured in an aqueous PBS buffer to calibrate geometrical parameters (lateral radius, *w*_0_, and aspect ratio, *s*) of the confocal detection volumes, which were approximated as 3D Gaussians.

For exocyst proteins diffusing in live cells, a 2-component diffusion model was applied to fit the three correlation curves, namely green auto-correlation (for GFP), red auto-correlation (for JF-646), and cross-correlation:

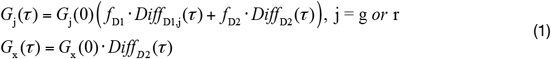

where the diffusion terms are expressed as:

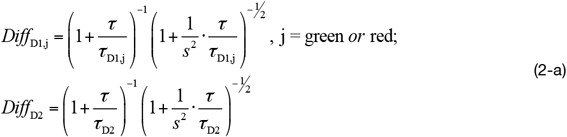

for 3-dimensional diffusion in the cytosol, or:

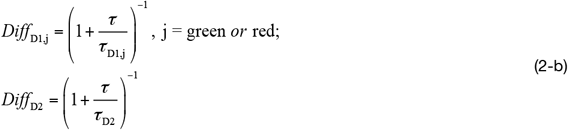

for 2-dimensional diffusion at the membrane.

Note that the cross-correlation contains only one diffusion component, which has the same parameters as the slow diffusion component of the green and the red auto-correlation curves. In Eqs. 2, τ_D1_ and τ_D2_ are the residence times of the two diffusing species, and *f*_D1_ and *f*_D2_ are corresponding fluorescence intensity contributions of the two diffusion species (*f*_D1_ + *f*_D2_ = 1). In the case of monomeric proteins, *f*_D1_ and *f*_D2_ equal to the concentration fractions of the two diffusing species. Diffusion coefficients (*D*) were calculated from fitted estimate of *τ*_D_ values as 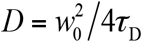, where *w*_0_ is the lateral radius of the detection volume.

Fractions of coupling are then calculated from the correlation amplitudes after fitting to Eqs. 1^51^:

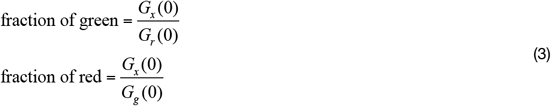

Fractions of coupling calculated from Eqs. 3 were corrected for spectral cross-talk using available fluorescence intensity information^52^. Correction factors were consistently lower than 0.1% owing to the high fluorescence intensity of the JF-646 dye relative to GFP. In addition, the fractions were not corrected for possible detection volume misoverlap due to the lack of a reliable in-cell control sample. This may lead to an underestimation by a constant scaling factor, which does not affect the comparison of measured fractions of coupling across different cells/samples.

The hydrodynamic radius (*R_H_*) of the fluorescent particles measured by FCS can be estimated from the diffusion coefficient using the Stokes-Einstein equation:

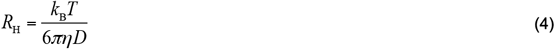

where *k*_B_ is the Boltzmann constant, *T* is the measurement temperature (20°C), *η* is the viscosity of the solvent (viscosity of water was use for the calculation), and *D* is the diffusion coefficient estimated from the autocorrelation curve.

### Statistical Analysis

Data are reported as ± *s.d., ± s.e.m* or 25^th^-75^th^ IQR, and analyzed by Student’s t-test or oneway analysis of variance (ANOVA) as indicated using Graphpad Prism 7, RStudio, or JASP (ver.0.8.6) statistical softwares. For curve fitting, fits were assessed by accounting for variation in the FCS data curve and Chi-square test statistics. When using ANOVA or Kruskal-Wallis test, *post hoc* analysis was done using Tukey’s, Dunn’s or Scheffe’s multiple comparison tests as appropriate. All statistical analysis was considered significant at *p* < 0.05.

**Supplementary Table 1.**
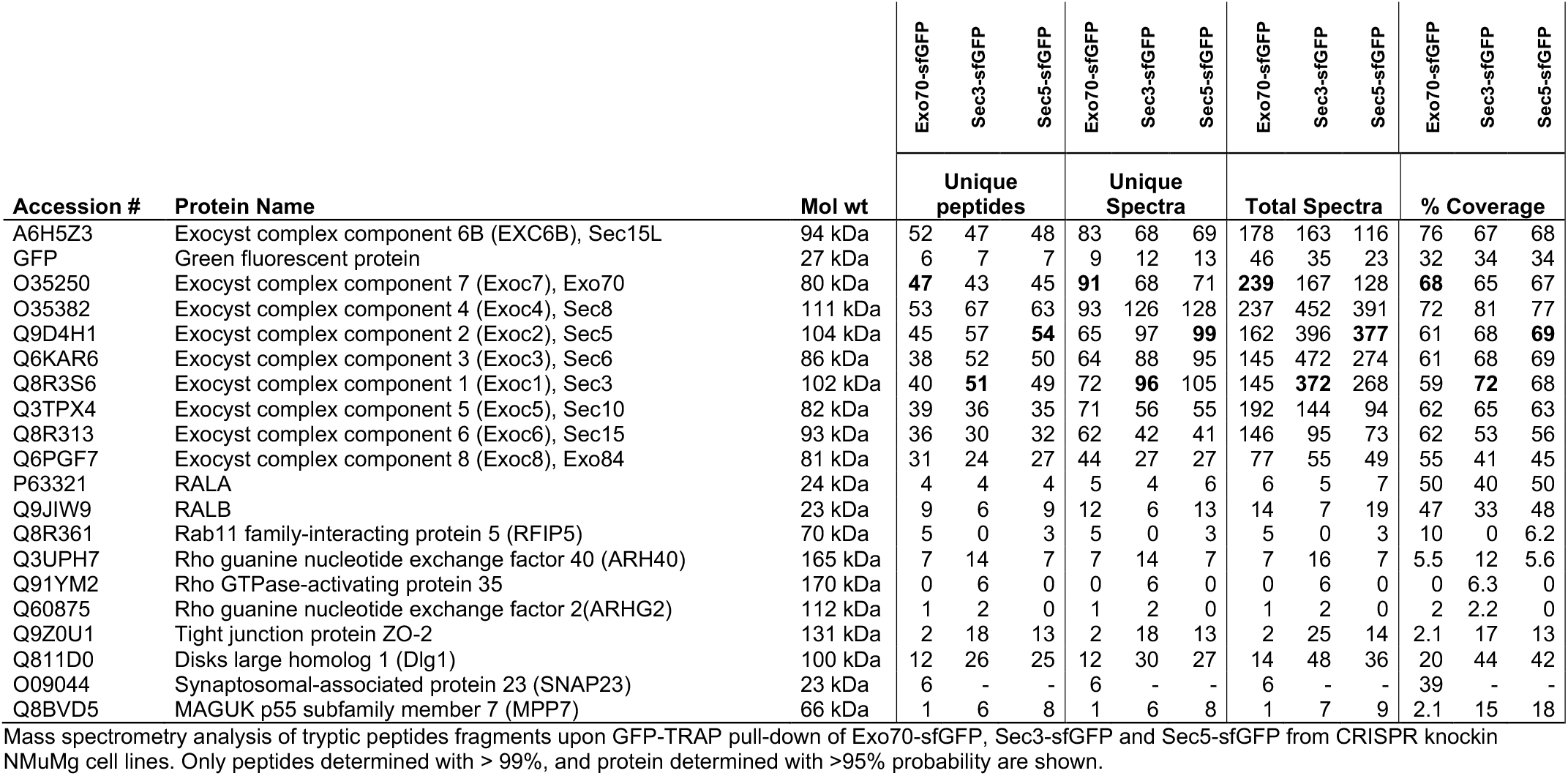
LC-MS/MS analysis of exocyst subunits interactions

**Supplementary Table 2.**
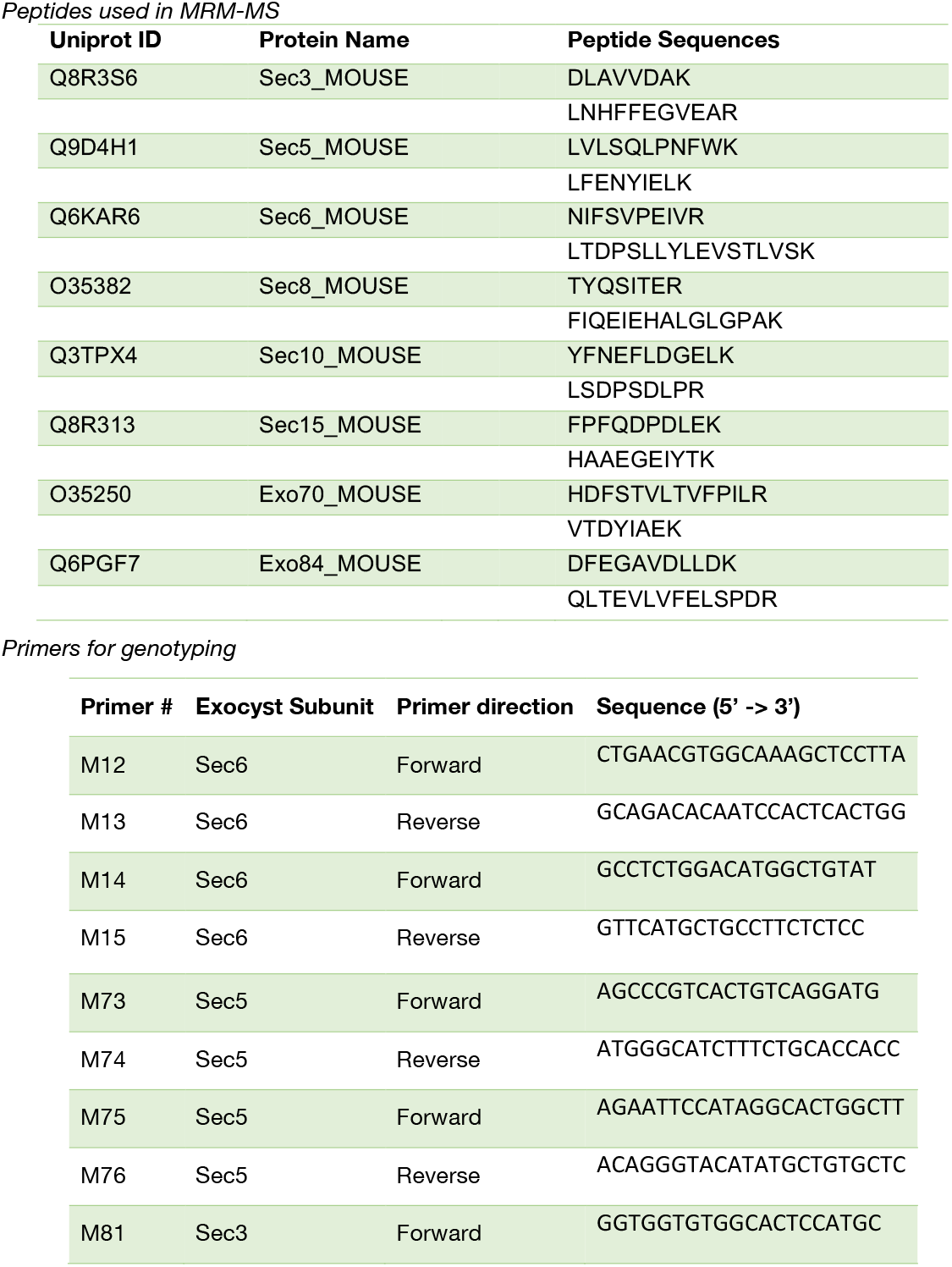
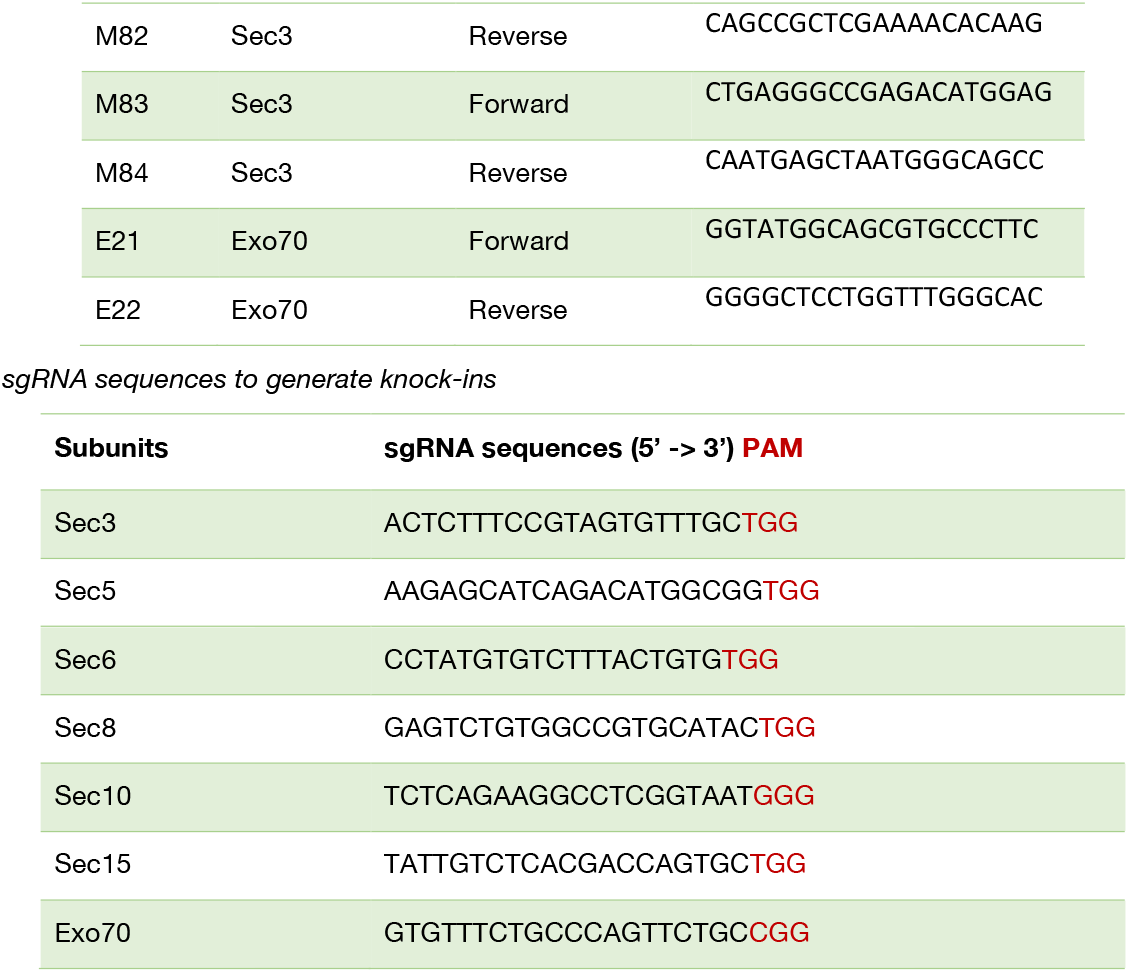
Peptides and primers sequences.

## References

1. Dubuke, M.L. & Munson, M. The Secret Life of Tethers: The Role of Tethering Factors in SNARE Complex Regulation. Front Cell Dev Biol 4, 42 (2016).

2. Brunet, S. & Sacher, M. Are all multisubunit tethering complexes bona fide tethers? Traffic 15, 1282–1287 (2014).

3. Chou, H.T., Dukovski, D., Chambers, M.G., Reinisch, K.M. & Walz, T. CATCHR, HOPS and CORVET tethering complexes share a similar architecture. Nat Struct Mol Biol 23, 761–763 (2016).

4. Fotso, P., Koryakina, Y., Pavliv, O., Tsiomenko, A.B. & Lupashin, V.V. Cog1p plays a central role in the organization of the yeast conserved oligomeric Golgi complex. J Biol Chem 280, 27613–27623 (2005).

5. Ungar, D., Oka, T., Vasile, E., Krieger, M. & Hughson, F.M. Subunit architecture of the conserved oligomeric Golgi complex. J Biol Chem 280, 32729–32735 (2005).

6. Willett, R. et al. COG complexes form spatial landmarks for distinct SNARE complexes. Nature Commun 4, 1553 (2013).

7. Heider, M.R. & Munson, M. Exorcising the exocyst complex. Traffic (Copenhagen, Denmark) 13, 898–907 (2012).

8. Wu, B. & Guo, W. The Exocyst at a Glance. J Cell Sci 128, 2957–2964 (2015).

9. TerBush, D.R., Maurice, T., Roth, D. & Novick, P. The Exocyst is a multiprotein complex required for exocytosis in Saccharomyces cerevisiae. EMBO J 15, 6483–6494 (1996).

10. Guo, W., Grant, A. & Novick, P. Exo84p is an exocyst protein essential for secretion. J Biol Chem 274, 23558–23564 (1999).

11. Zhang, X. et al. Membrane association and functional regulation of Sec3 by phospholipids and Cdc42. J Cell Biol 180, 145–158 (2008).

12. He, B., Xi, F., Zhang, X., Zhang, J. & Guo, W. Exo70 interacts with phospholipids and mediates the targeting of the exocyst to the plasma membrane. EMBO J 26, 4053–4065 (2007).

13. Yakir-Tamang, L. & Gerst, J.E. A phosphatidylinositol-transfer protein and phosphatidylinositol-4-phosphate 5-kinase control Cdc42 to regulate the actin cytoskeleton and secretory pathway in yeast. Mol Biol Cell 20, 3583–3597 (2009).

14. Baek, K. et al. Structure-function study of the N-terminal domain of exocyst subunit Sec3. J Biol Chem 285, 10424–10433 (2010).

15. Luo, G., Zhang, J. & Guo, W. The role of Sec3p in secretory vesicle targeting and exocyst complex assembly. Mol Biol Cell 25, 3813–3822 (2014).

16. Liu, D. & Novick, P. Bem1p contributes to secretory pathway polarization through a direct interaction with Exo70p. J Cell Biol 207, 59–72 (2014).

17. Liu, D., Li, X., Shen, D. & Novick, P. Two subunits of the exocyst, Sec3p and Exo70p, can function exclusively on the plasma membrane. Mol Biol Cell 29, 736–750 (2018).

18. Boyd, C., Hughes, T., Pypaert, M. & Novick, P. Vesicles carry most exocyst subunits to exocytic sites marked by the remaining two subunits, Sec3p and Exo70p. J Cell Biol 167, 889–901 (2004).

19. Heider, M.R. et al. Subunit connectivity, assembly determinants and architecture of the yeast exocyst complex. Nat Struct Mol Biol 23, 59–66 (2016).

20. Donovan, K.W. & Bretscher, A. Myosin-V is activated by binding secretory cargo and released in coordination with Rab/exocyst function. Dev cell 23, 769–781 (2012).

21. Donovan, K.W. & Bretscher, A. Tracking individual secretory vesicles during exocytosis reveals an ordered and regulated process. J Cell Biol 210, 181–189 (2015).

22. Moskalenko, S. et al. Ral GTPases regulate exocyst assembly through dual subunit interactions. J Biol Chem 278, 51743–51748 (2003).

23. Yamashita, M. et al. Structural basis for the Rho-and phosphoinositide-dependent localization of the exocyst subunit Sec3. Nat Struct Mol Biol 17, 180–186 (2010).

24. Wu, H., Turner, C., Gardner, J., Temple, B. & Brennwald, P. The Exo70 subunit of the exocyst is an effector for both Cdc42 and Rho3 function in polarized exocytosis. Mol Biol Cell 21, 430–442 (2010).

25. Adamo, J.E., Rossi, G. & Brennwald, P. The Rho GTPase Rho3 has a direct role in exocytosis that is distinct from its role in actin polarity. Mol Biol Cell 10, 4121–4133 (1999).

26. Pathak, R. et al. The microtubule-associated Rho activating factor GEF-H1 interacts with exocyst complex to regulate vesicle traffic. Dev cell 23, 397–411 (2012).

27. Das, A. & Guo, W. Rabs and the exocyst in ciliogenesis, tubulogenesis and beyond. Trends Cell Biol 21, 383–386 (2011).

28. Guo, W., Tamanoi, F. & Novick, P. Spatial regulation of the exocyst complex by Rho1 GTPase. Nat Cell Biol 3, 353–360 (2001).

29. Zhang, X. et al. Cdc42 interacts with the exocyst and regulates polarized secretion. The J Biol Chem 276, 46745–46750 (2001).

30. Ahmed, S.M. & Macara, I.G. The Par3 polarity protein is an exocyst receptor essential for mammary cell survival. Nature Commun 8, 14867 (2017).

31. Rivera-Molina, F. & Toomre, D. Live-cell imaging of exocyst links its spatiotemporal dynamics to various stages of vesicle fusion. J Cell Biol 201, 673–680 (2013).

32. Friedrich, G.A., Hildebrand, J.D. & Soriano, P. The secretory protein Sec8 is required for paraxial mesoderm formation in the mouse. Dev Biol 192, 364–374 (1997).

33. Murthy, M. et al. Sec6 mutations and the Drosophila exocyst complex. J Cell Sci 118, 1139–1150 (2005).

34. Hutagalung, A.H., Coleman, J., Pypaert, M. & Novick, P.J. An internal domain of Exo70p is required for actin-independent localization and mediates assembly of specific exocyst components. Mol Biol Cell 20, 153–163 (2009).

35. Roumanie, O. et al. Rho GTPase regulation of exocytosis in yeast is independent of GTPase hydrolysis and polarization of the exocyst complex. J Cell Biol 170, 583–594 (2005).

36. Torres, M.J. et al. Role of the Exocyst Complex Component Sec6/8 in Genomic Stability. Mol Cell Biol 35, 3633–3645 (2015).

37. Chen, X.W. et al. Exocyst function is regulated by effector phosphorylation. Nat Cell Biol 13, 580–588 (2011).

38. Balakireva, M. et al. The Ral/exocyst effector complex counters c-Jun N-terminal kinase-dependent apoptosis in Drosophila melanogaster. Mol Cell Biol 26, 8953–8963 (2006).

39. Yeaman, C., Grindstaff, K.K. & Nelson, W.J. Mechanism of recruiting Sec6/8 (exocyst) complex to the apical junctional complex during polarization of epithelial cells. J Cell Sci 117, 559–570 (2004).

40. Mei, K. et al. Cryo-EM structure of the exocyst complex. Nat Struct Mol Biol 25, 139–146 (2018).

41. Kreitzer, G. et al. Three-dimensional analysis of post-Golgi carrier exocytosis in epithelial cells. Nature Cell Biol 5, 126–136 (2003).

42. Gura Sadovsky, R., Brielle, S., Kaganovich, D. & England, J.L. Measurement of Rapid Protein Diffusion in the Cytoplasm by Photo-Converted Intensity Profile Expansion. Cell Rep 18, 2795–2806 (2017).

43. Wu, J.Q., McCormick, C.D. & Pollard, T.D. Chapter 9: Counting proteins in living cells by quantitative fluorescence microscopy with internal standards. Methods Cell Biol 89, 253–273 (2008).

44. Latty, S.L. et al. Referenced Single-Molecule Measurements Differentiate between GPCR Oligomerization States. Biophys J 109, 1798–1806 (2015).

45. Picco, A. et al. The In Vivo Architecture of the Exocyst Provides Structural Basis for Exocytosis. Cell 168, 400–412 e418 (2017).

46. Ahmed, S.M. & Macara, I.G. The Par3 polarity protein is an exocyst receptor essential for mammary cell survival. Nature Commun (2017).

47. Grimm, J.B. et al. A general method to fine-tune fluorophores for live-cell and in vivo imaging. Nat Methods 14, 987–994 (2017).

48. Shivnaraine, R.V. et al. Single-Molecule Analysis of the Supramolecular Organization of the M2 Muscarinic Receptor and the Galphai1 Protein. J Am Chem Soc 138, 11583–11598 (2016).

49. James, J.R., Oliveira, M.I., Carmo, A.M., Iaboni, A. & Davis, S.J. A rigorous experimental framework for detecting protein oligomerization using bioluminescence resonance energy transfer. Nat Methods 3, 1001–1006 (2006).

50. Zhang, Z., Yomo, D. & Gradinaru, C. Choosing the right fluorophore for single-molecule fluorescence studies in a lipid environment. Biochim Biophys Acta 1859, 1242–1253 (2017).

51. Bacia, K. & Schwille, P. Practical guidelines for dual-color fluorescence cross-correlation spectroscopy. Nat Protoc 2, 2842–2856 (2007).

52. Bacia, K., Petrasek, Z. & Schwille, P. Correcting for spectral cross-talk in dual-color fluorescence cross-correlation spectroscopy. Chemphyschem 13, 1221–1231 (2012).

